# Metformin inhibits mitochondrial complex I in intestinal epithelium to promote glycemic control

**DOI:** 10.1101/2025.10.02.678294

**Authors:** Zachary L. Sebo, Ram P. Chakrabarty, Rogan A. Grant, Karis B. D’Alessandro, Alec R. Koss, Jenna L. E. Blum, Shawn M. Davidson, Colleen R. Reczek, Navdeep S. Chandel

**Affiliations:** Department of Medicine, Division of Pulmonary and Critical Care Medicine, Northwestern University Feinberg School of Medicine, Chicago, IL, USA; Department of Biochemistry and Molecular Genetics, Northwestern University Feinberg School of Medicine, Chicago, IL, USA; Chan Zuckerberg Biohub, Chicago, IL, USA

## Abstract

Metformin is a therapeutically versatile biguanide drug primarily prescribed for type II diabetes. Despite its extensive use, the mechanisms underlying many of its clinical effects, including attenuated postprandial glucose excursions, elevated intestinal glucose uptake, and increased production of lactate, Lac-Phe and GDF15, remain unclear. Here, we map these and other clinical effects of metformin to intestine-specific mitochondrial complex I inhibition. Using human metabolomic data and an orthogonal genetics approach in male mice, we demonstrate that metformin suppresses citrulline synthesis, a metabolite generated exclusively by small intestine mitochondria, and increases GDF15 by inhibiting the mitochondrial respiratory chain at complex I. This inhibition co-opts the intestines to function as a glucose sink, driving uptake of excess glucose and converting it to lactate and Lac-Phe. Notably, the glucose-lowering effect of another biguanide, phenformin, and berberine, a structurally unrelated nutraceutical, similarly depends on intestine-specific mitochondrial complex I inhibition, underscoring a shared therapeutic mechanism.

## Introduction

Metformin is the most widely prescribed medication for type II diabetes and the only FDA-approved drug of the biguanide class^1,2^. Prior to the 1990s, metformin was thought to promote glycemic control primarily by enhancing glucose utilization in peripheral tissues, including the intestines, with inhibition of endogenous glucose production considered secondary^3^. However, due to technological advances in isotope tracing and NMR, subsequent studies demonstrated a suppressive effect of metformin on endogenous glucose production^4–6^, which is now generally attributed to the direct inhibition of hepatic gluconeogenesis^2,7^. Metformin was initially proposed to elicit its anti-gluconeogenic effect by inhibiting mitochondrial complex I of the electron transport chain in hepatocytes^8,9^. However, this explanation has been refuted because inhibiting complex I requires millimolar concentrations of metformin, which are only attained in the intestines of patients on standard dosing regimens^10–13^. Consequently, alternative molecular targets within the liver have been put forward to explain how metformin suppresses gluconeogenesis^14–16^. In contrast, other studies have challenged the centrality of the liver in metformin’s mechanism of action, demonstrating that metformin does not reduce endogenous glucose production in patients with prediabetes or those with recent-onset or well-controlled type II diabetes^17–20^. Instead, the primary antidiabetic effect of metformin in these patient groups is enhanced glucose clearance^17,18^, concomitant with elevated aerobic glycolysis^17^. Indeed, metformin has consistently been shown to enhance glucose utilization in humans^21–26^.

Mostly due to advances in clinical imaging techniques, it is now clear that the intestines are the primary site of increased glycolytic activity, where metformin promotes both glucose uptake (as measured by 18F-fluorodeoxyglucose [FDG] accumulation) and lactate production^11,24,27–30^. 18F-fluorodeoxyglucose positron emission tomography (FDG-PET), a widely used imaging method for cancer detection, relies on the elevated uptake of FDG in tumors relative to normal tissues. Because metformin increases FDG accumulation in the small and large intestines, it can obscure tumors during imaging. Consequently, by the early 2010s, discontinuing metformin before FDG-PET scans became standard of care to avoid compromising cancer detection^31–33^.

Despite this clinical progress, both the relative contribution of the intestinal effects of metformin on glycemic control and the underlying mechanism remain unclear. To address this gap in understanding, a bona fide molecular target of metformin in the intestines must, at minimum, explain the drug’s ability to enhance intestinal glucose utilization and increase blood glucose clearance. Ideally, the inhibition of this molecular target would also account for multiple other clinical effects of metformin.

In this study, we leverage publicly available metabolomic data from humans and genetic tools in mice to pinpoint mitochondrial complex I as an essential therapeutic target of metformin in the intestinal epithelium. In addition to enhanced intestinal glucose utilization and blood glucose clearance, this mechanism accounts for metformin-induced citrulline depletion, improved postprandial glycemia, and elevated Lac-Phe and GDF15 levels—all of which are definitive clinical outcomes caused by metformin treatment. We further determined that phenformin, another biguanide, and berberine, a natural compound used as an over-the-counter treatment for type II diabetes, lower blood glucose through the same mechanism. Thus, we identify mitochondrial complex I in intestinal epithelium as a shared and essential therapeutic target for metformin, phenformin, and berberine.

## Results

### Metformin suppresses intestinal citrulline synthesis by inhibiting mitochondrial complex I

Postprandial glucose, which is elevated in type II diabetes, is a stronger predictor of cardiovascular disease and all-cause mortality than fasting glucose^34,35^. Metformin acutely suppresses these meal-induced glucose spikes^36–39^ and improves glucose tolerance, even in normoglycemic individuals^40^. We confirmed the acute glucose-lowering effect of metformin by analyzing publicly available data from a cohort of non-diabetic subjects, each of whom also underwent plasma metabolomic profiling (**Extended Data Fig. 1a-c**)^41^. By examining the metabolomics data, we found that citrulline was the most significantly downregulated metabolite by metformin (**Fig. 1a-c, Extended Data Table 1**)^41^. Similarly, in a separate cohort of obese patients with type II diabetes, treatment with metformin (1,500 mg per day for at least six months) led to a significant reduction in circulating citrulline levels compared to patients who did not receive metformin. (**Fig. 1d**)^42^. Consistent with these findings, other groups have reported a pronounced decrease in the citrulline levels of patients with type II diabetes receiving metformin^43,44^. Interestingly, the enzyme responsible for citrulline synthesis, ornithine transcarbamylase (OTC), is localized to the mitochondrial matrix and is exclusively expressed in the liver and small intestine. While liver-produced citrulline is locally metabolized to argininosuccinate as part of the urea cycle, intestine-produced citrulline is released into the circulation to augment systemic nitric oxide synthesis^45–48^. The penultimate enzyme in the two-step citrulline synthesis pathway, mitochondrial carbamoyl phosphate synthetase I (CPS1), has the same expression pattern and subcellular localization as OTC and requires mitochondrial ATP, rather than ATP produced by glycolysis, to function (**Extended Data Fig. 3a, b**). Importantly, ATP generated in the cytosol from glycolysis does not contribute to the ATP pool in the mitochondrial matrix^49,50^ and therefore cannot compensate for reduced oxidative phosphorylation to sustain citrulline synthesis. Indeed, patients with mitochondrial dysfunction due to a mutation in *LRPPRC* display reduced plasma citrulline levels^51^. Thus, it is plausible that metformin decreases citrulline by suppressing mitochondrial activity in the small intestine.

**Figure 1.**
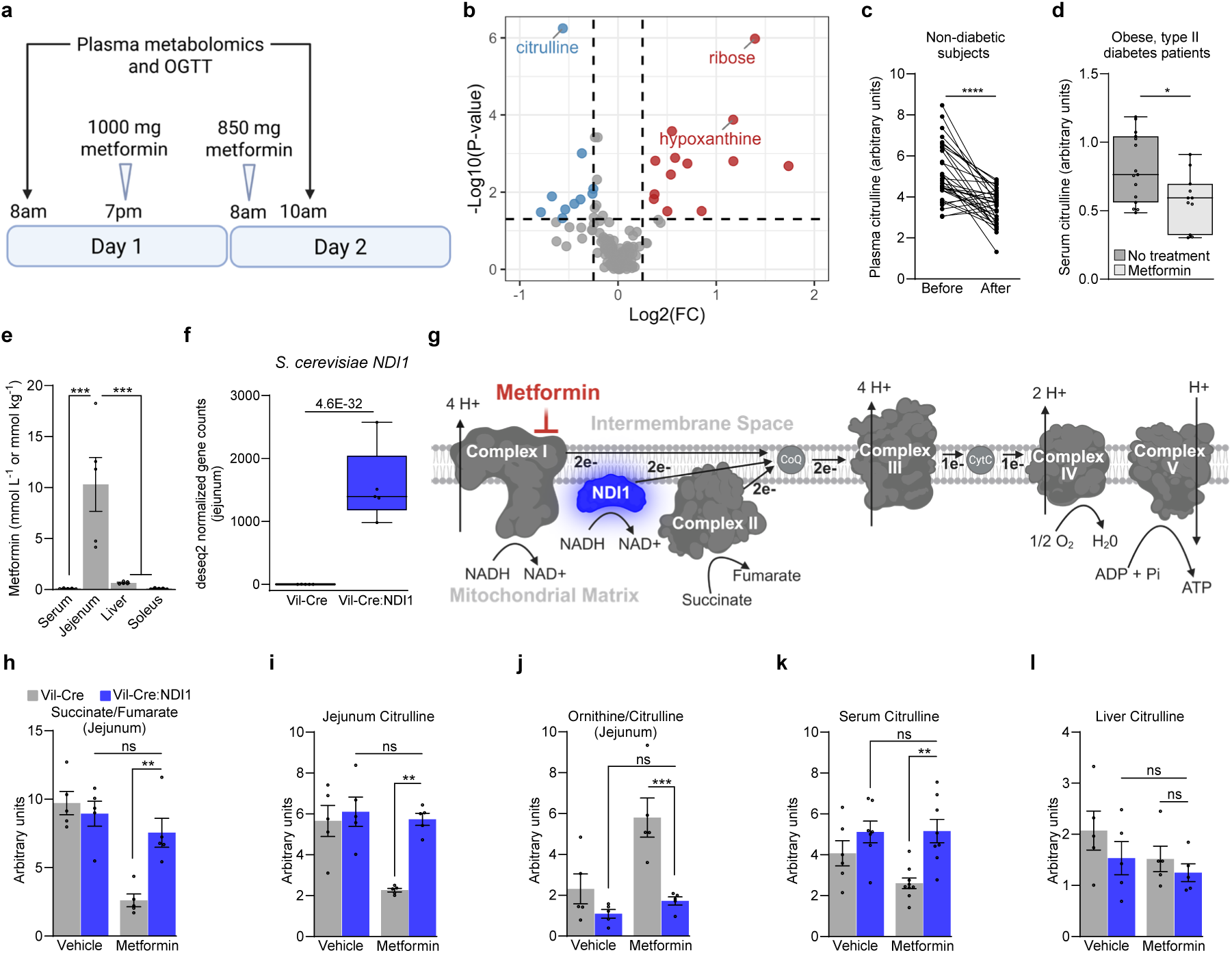
Metformin inhibits mitochondrial complex I in intestinal epithelium to suppress citrulline synthesis. (**a**) Schematic illustrating sample collection and metformin dosing in the Rotroff et al. cohort; related to panels (b) and (c). Created with BioRender.com. (**b**) Volcano plot of 155 known metabolites in blood plasma before and after metformin from the Rotroff et al. cohort of human study participants; metabolites downregulated by metformin are in blue; metabolites upregulated by metformin are in red; the top 3 metabolites are labeled. n=33. (**c**) Plasma citrulline levels in the Rotroff et al. cohort before and after metformin. n=33. (**d**) Serum citrulline levels in the Aleidi et al. cohort of patients with obesity and type II diabetes. n=15 no treatment, n=11 metformin. (**e**) Metformin concentrations in serum and different organs of overnight fasted C57BL/6J mice one hour after a single oral dose of metformin (200 mg kg^-1^). n=5. (**f**) *Ndi1* normalized gene expression in the jejunum of Vil-Cre and Vil-Cre:NDI1 mice. n=5 per condition (q = 4.6 x 10^-32^, Wald test). (**g**) Schematic of the mitochondrial electron transport chain and NDI1. Created with BioRender.com. (**h**) Succinate/fumarate ratio of jejunum in overnight fasted mice one hour after oral administration of vehicle or metformin (200 mg kg^-1^). n=5 per condition. (**i**) Jejunum citrulline levels normalized to total ion count one hour after orally administered vehicle (water) or metformin (200 mg kg^-1^) in overnight fasted mice. n=5 per condition. (**j**) Ornithine/Citrulline ratio in jejunum one hour after oral administration of vehicle (water) or metformin (200 mg kg^-1^) in overnight-fasted mice; n=5 per condition. (**k**) Serum citrulline levels 2.5h after oral administration of vehicle (water) or metformin (200 mg kg^-1^) in overnight fasted mice (normalized to thymine-D4 internal standard). Mice were also given an oral bolus of glucose (2 g kg^-1^) 30 minutes after vehicle/metformin administration; Vil-Cre^vehicle^ n=6, Vil-Cre:NDI1^vehicle^ n=7, Vil-Cre^metformin^ n=9, Vil-Cre:NDI1^metformin^ n=8. (**l**) Liver citrulline levels normalized to total ion count one hour after orally administered vehicle (water) or metformin (200 mg kg^-1^) in overnight fasted mice. n=5 per condition. OGTT = oral glucose tolerance test. All mice were males and 8-12 weeks of age. The vehicle was water in all experiments. For (e and h-l), results represent mean ± SEM. For (d, f), results are presented as min-to-max box and whisker plots. Statistical significance for (c) was determined by Paired t test. Statistical significance for (d) was determined by Unpaired t test. For (e), One-way ANOVA with Bonferroni correction for multiple comparisons. For (h-l), Two-way ANOVA with Bonferroni correction for multiple comparisons. *P<0.05, **P<0.01, ***P<0.001, ****P<0.0001.

In line with this concept, intestinal metformin levels exceed those in plasma by up to 300-fold and those in the liver by 10-100 fold, with local concentrations reaching well into the millimolar range (**Fig. 1e**)^11–13,52^. This is critical, as in vitro and structural studies show that metformin inhibits mitochondrial complex I only at concentrations typically achieved in the intestines under standard clinical dosing^8,9,53,54^. Mitochondrial complex I is a large (∼1 megadalton) protein assembly of 45 subunits and is embedded in the inner mitochondrial membrane where it supports oxidative phosphorylation and TCA cycle activity by donating electrons to ubiquinone, regenerating NAD^+^, and pumping protons into the inner membrane space. Mitochondrial complex I can also generate superoxide through reverse electron transport^55^. Biguanides like metformin require an intact inner mitochondrial membrane potential to accumulate in the matrix^8,56^ and reversibly inhibit mitochondrial complex I by binding essential residues within the ubiquinone binding channel, stabilizing the enzyme’s deactive form^54^.

Our group recently reported direct evidence that mitochondrial complex I inhibition is necessary for metformin to lower blood glucose in vivo^57^. By generating transgenic mice that ubiquitously express *Saccharomyces cerevisiae* NADH dehydrogenase (NDI1), we conferred resistance to mitochondrial complex I inhibition in all tissues of the body. NDI1 localizes to the inner mitochondrial membrane where it acts as a homodimer to catalyze ubiquinone reduction coupled to NAD^+^ regeneration, without pumping protons or generating superoxide^58,59^ (**Fig. 1g**). NDI1 rescues genetic complex I dysfunction and is insensitive to pharmacologic mitochondrial complex I inhibitors including biguanides^56,60–62^. Therefore, NDI1 can be expressed in mammalian systems to maintain electron transport chain activity in a manner that bypasses mitochondrial complex I. Mice with ubiquitous NDI1 expression were partially resistant to the glucose-lowering effect of metformin, demonstrating that complex I inhibition contributes to the antidiabetic effect of this drug^57^. Nevertheless, the specific tissue in which mitochondrial complex I is inhibited, as well as the downstream mechanism were not identified.

In the present study, we generated mice that specifically express NDI1 in intestinal epithelial cells by crossing Villin-Cre mice to animals that harbor NDI1 with a lox-stop-lox sequence targeted to the ROSA26 locus^61^ (Vil-Cre:NDI1 mice) (**Fig. 1f, g**). RNA-sequencing analysis revealed a negligible effect of NDI1 on the intestinal transcriptome and NDI1 was not expressed in the liver of Vil-Cre:NDI1 mice (**Extended Data Fig. 2a, b).** When mitochondrial complex I is inhibited, complex II (succinate dehydrogenase) becomes the primary entry point for electrons into the respiratory chain^63^. Under these conditions, forward flux through succinate dehydrogenase is elevated, resulting in the rapid consumption of succinate and production of fumarate, causing a decrease in the succinate/fumarate ratio^64^. Metformin decreases the intestinal succinate/fumarate ratio in control animals, indicating increased flux through succinate dehydrogenase. However, the succinate/fumarate ratio was unchanged by metformin in mice with intestinal NDI1 expression (**Fig. 1h**). Thus, metformin inhibits mitochondrial complex I in intestinal epithelium in vivo. We further assessed metformin’s effect on intestinal tissue by profiling the relative abundance of ∼250 hydrophilic metabolites in the jejunum by liquid chromatography-mass spectrometry (LCMS). Hierarchical clustering of metabolomic data showed no effect of NDI1 in vehicle-treated animals. However, NDI1 substantially diminished metformin-induced changes to the intestinal metabolome (**Extended Data Fig. 4a**). Differentially abundant metabolites were identified between metformin-treated animals with and without NDI1 (**Extended Data Table 2**). We found that citrulline showed the most significant decrease among all metabolites in response to metformin, and this reduction depended on inhibition of mitochondrial complex I (**Fig. 1i, Extended Data Fig. 4b**). Similarly, we observed a marked decrease in circulating citrulline levels, along with an increased ornithine/citrulline ratio in the jejunum, indicating reduced flux through OTC (**Fig. 1j, k**). In contrast, no change in liver citrulline was observed (**Fig. 1l**). Together, these data demonstrate that metformin suppresses intestinal citrulline synthesis by inhibiting mitochondrial complex I.

### Metformin inhibits mitochondrial complex I to drive intestinal glucose disposal

Inhibition of mitochondrial complex I or other components of the electron transport chain typically results in a compensatory increase in glycolysis^65^. Indeed, an underappreciated clinical effect of metformin is enhanced intestinal glucose uptake, as measured by ^18^F-fluorodeoxyglucose positron emission tomography (FDG-PET)^27–29,33^. FDG is a radioactive glucose mimetic and measuring its uptake is a standard clinical cancer detection method due to the enhanced glucose uptake of tumors^66^. Discontinuation of metformin before FDG-PET scans is standard of care because metformin confounds image interpretation by enhancing intestinal FDG accumulation^31–33^. However, the underlying mechanism of this clinical effect is unknown. We hypothesized that metformin targets mitochondrial complex I to drive intestinal glucose uptake and glycolysis (**Fig. 2a**). To test this, we performed FDG-PET on Vil-Cre:NDI1 mice treated with metformin. Consistent with clinical observations, metformin elevates intestinal FDG uptake in control mice (**Fig. 2b, c**). However, in Vil-Cre:NDI1 mice, metformin fails to induce FDG uptake in the intestines (**Fig. 2b, c**). Importantly, metformin did not alter FDG accumulation in the liver or muscle in either genotype, indicating that its effect is specific to the intestines (**Extended Data Fig. 5a, b**). We also performed an orthogonal glucose-uptake assay by intraperitoneally injecting mice with 2-deoxyglucose (2DG), another glucose mimetic. 2DG is taken up by cells and converted to 2-deoxyglucose-6-phosphate (2DG6P) but cannot be further metabolized. Metformin increases intestinal 2DG6P accumulation only in mice without NDI1 expression (**Fig. 2d**). In addition, we assessed the mRNA expression of intestinal glucose transporters and found no differences in response to metformin or NDI1 (**Extended Data Fig. 5c-f**). Thus, mitochondrial complex I inhibition is necessary for metformin to drive intestinal glucose uptake in a manner that does not involve the transcriptional upregulation of glucose transporters.

**Figure 2.**
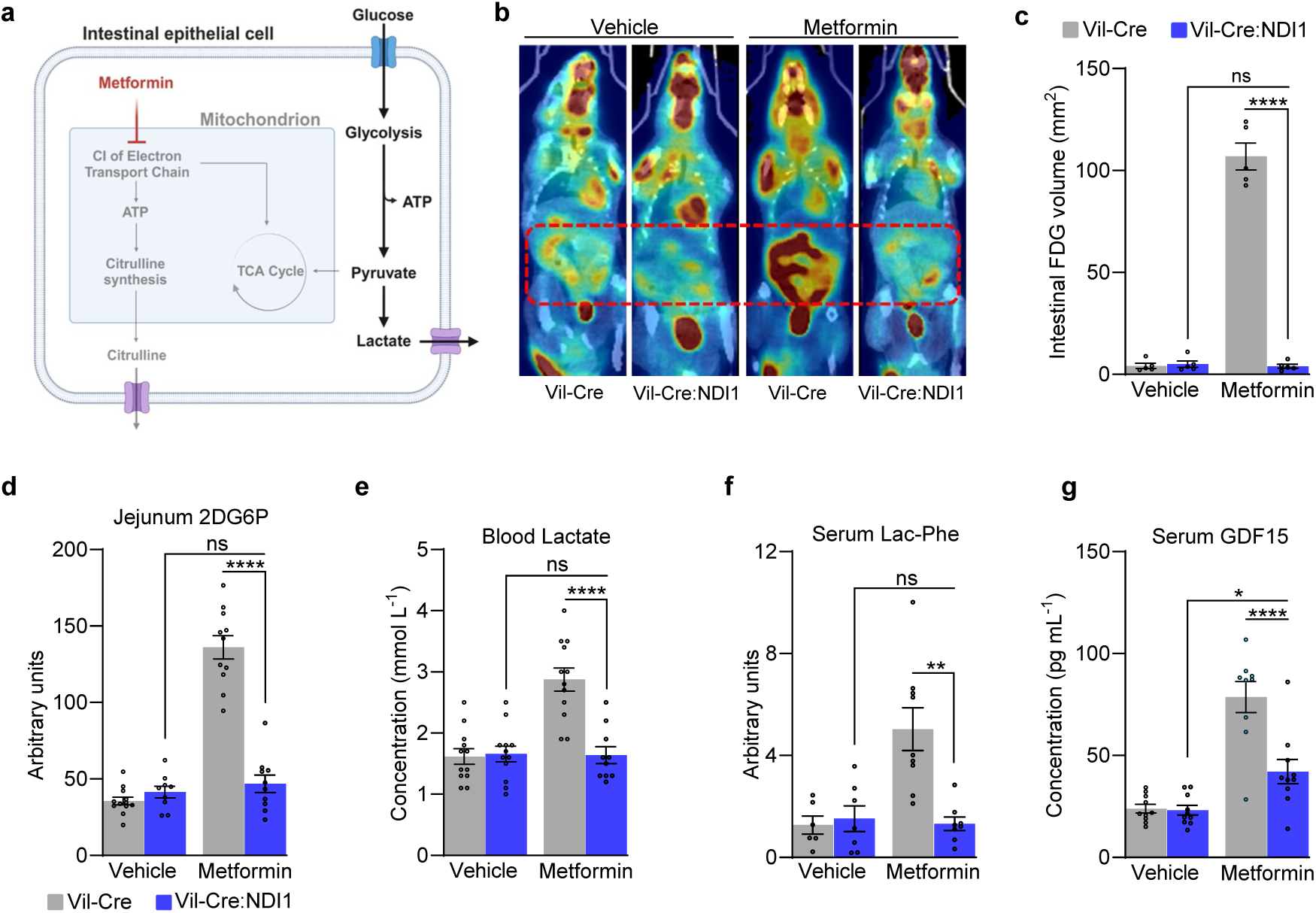
Metformin inhibits mitochondrial complex I in intestinal epithelium to drive glucose uptake and glycolysis. (**a**) Model of metformin-mediated glucose disposal and suppressed citrulline synthesis in intestinal epithelium. Created with BioRender.com. (**b**) Representative FDG-PET images of Vil-Cre and Vil-Cre:NDI1 mice with and without metformin. Mice were fasted overnight, followed by an oral gavage of vehicle or metformin (200 mg kg^-1^); 30 minutes later, FDG was administered via tail vein injection; PET imaging began 40 minutes after FDG administration and completed 60 minutes after FDG administration. (**c**) Volume quantification of FDG signal in mice treated with vehicle or 200 mg kg^-1^ metformin; SUV threshold of 0.45, n=5 per condition. (**d**) Relative 2-deoxyglucose-6-phosphate (2DG6P) in the jejunum of Vil-Cre and Vil-Cre:NDI1 mice. Mice were fasted overnight, followed by an oral gavage of vehicle or metformin (200 mg kg^-1^); 30 minutes later, 2-deoxyglucose (50 mg kg^-1^) was administered in combination with 2 g kg^-1^ glucose via intraperitoneal injection; one hour later jejunum was harvested to measure 2DG6P. Vil-Cre^vehicle^ n=12, Vil-Cre:NDI1^vehicle^ n=9, Vil-Cre^metformin^ n=11, Vil-Cre:NDI1^metformin^ n=10. (**e**) Blood lactate levels in overnight fasted mice. Mice were orally administered vehicle (water) or metformin (200 mg kg^-1^), followed by an oral gavage of glucose (2 g kg^-1^) 30 minutes later; 30 minutes after glucose administration, blood lactate was measured; Vil-Cre^vehicle^ n=12, Vil-Cre:NDI1^vehicle^ n=12, Vil-Cre^metformin^ n=12, Vil-Cre:NDI1^metformin^ n=10. (**f**) Serum lactoyl-phenylalanine (Lac-Phe) levels 2.5h after oral administration of vehicle (water) or metformin (200 mg kg^-1^) in overnight fasted mice (normalized to thymine-D4 internal standard). Mice were also given an oral bolus of glucose (2 g kg^-1^) 30 minutes after vehicle/metformin administration; Vil-Cre^vehicle^ n=6, Vil-Cre:NDI1^vehicle^ n=7, Vil-Cre^metformin^ n=9, Vil-Cre:NDI1^metformin^ n=8. (**g**) Serum GDF15 levels in overnight fasted mice 8 hours after oral administration of vehicle or metformin (200 mg kg^-1^); Vil-Cre^vehicle^ n=10, Vil-Cre:NDI1^vehicle^ n=9, Vil-Cre^metformin^ n=9, Vil-Cre:NDI1^metformin^ n=10. All mice were males and 8-12 weeks of age. The vehicle was water for oral delivery and sterile PBS for injections. Statistical significance in (d-g) was determined by Two-way ANOVA with Bonferroni’s correction for multiple comparisons. **P<0.01, ***P<0.001, ****P<0.0001.

Lactate production is elevated in highly glycolytic cells, and metformin increases blood lactate in humans. Other biguanides, such as phenformin, were withdrawn from clinical use due to increased risk of lactic acidosis^67^. Thus, we hypothesized that metformin increases blood lactate by inhibiting mitochondrial complex I in intestinal epithelium (**Fig. 2a**). Consistent with this hypothesis, a single oral dose of metformin elevates blood lactate in control but not Vil-Cre:NDI1 mice (**Extended Data Fig. 5g**). Giving mice an oral bolus of glucose in addition to metformin further increases blood lactate in control mice. However, Vil-Cre:NDI1 mice remain insensitive to metformin-induced lactate production (**Fig. 2e**). Similar effects were observed with phenformin (**Extended Data Fig. 5h, i**). Metformin-induced glucose-to-lactate conversion was directly assessed by measuring the m+3 lactate / m+6 glucose ratio in blood serum after an oral dose of U-^13^C_6_-glucose. As expected, the m+3 lactate / m+6 glucose ratio was elevated by metformin in control but not Vil-Cre:NDI1 mice (**Extended Data Fig. 5j**). Together, these data demonstrate that mitochondrial complex I inhibition is required for biguanides to enhance glycolytic activity in intestinal epithelium.

Lactoyl-phenylalanine (Lac-Phe), a lactate derivative, and GDF15, a downstream effector of the mitochondrial integrated stress response, are also elevated by metformin in rodents and humans^68–73^. We found that Vil-Cre:NDI1 mice are refractory to metformin-induced Lac-Phe and GDF15 upregulation (**Fig. 2f, g**), indicating mitochondrial complex I inhibition in intestinal epithelium is necessary for metformin to elevate Lac-Phe and GDF15 levels. Metformin modestly lowers body weight in humans (∼3% loss over a year)^74^. Because Lac-Phe and GDF15 have been implicated in this effect^68–73^, we assessed body weight and food intake in diet-induced obese mice treated daily with metformin (200 mg kg^-1^) for two weeks. We found that metformin tended to slow weight gain but did not cause weight loss (**Extended Data Fig. 6b, c**), consistent with previous studies in mice which required higher doses of metformin (≥300 mg kg^-1^) to observe weight loss^68,70^.

### Intestine-specific inhibition of mitochondrial complex I is necessary for the antihyperglycemic effect of metformin

To determine whether intestinal mitochondrial complex I inhibition is necessary for the therapeutic glucose-lowering effect of metformin, we first performed glucose tolerance tests on mice fed a standard diet. Control and Vil-Cre:NDI1 mice have the same baseline glucose tolerance (**Fig. 3a-f**). However, the blood glucose-lowering effect of acutely administered metformin is significantly impaired in Vil-Cre:NDI1 mice (**Fig. 3a-f**). This occurs independently of the glucose administration route (oral gavage or intraperitoneal injection) (**Fig. 3c-f**) and at a low dose (100 mg kg^-1^) of metformin (**Fig. 3a, b**). Glucose tolerance was also assessed in diet-induced obese mice. While NDI1 did not affect body weight (**Extended Data Fig. 6a**), blood glucose lowering by metformin was attenuated in animals with intestinal NDI1 expression (**Fig. 4a-f**). Notably, resistance to metformin mediated by intestinal NDI1 was incomplete, consistent with our earlier findings in mice with ubiquitous NDI1 expression^57^, and tended to be greater at a low dose of metformin and when glucose was administered intraperitoneally rather than orally. This partial resistance suggests that metformin may engage therapeutic targets beyond mitochondrial complex I and could also reflect the inability of NDI1 to fully rescue metformin-induced suppression of complex I function.

**Figure 3.**
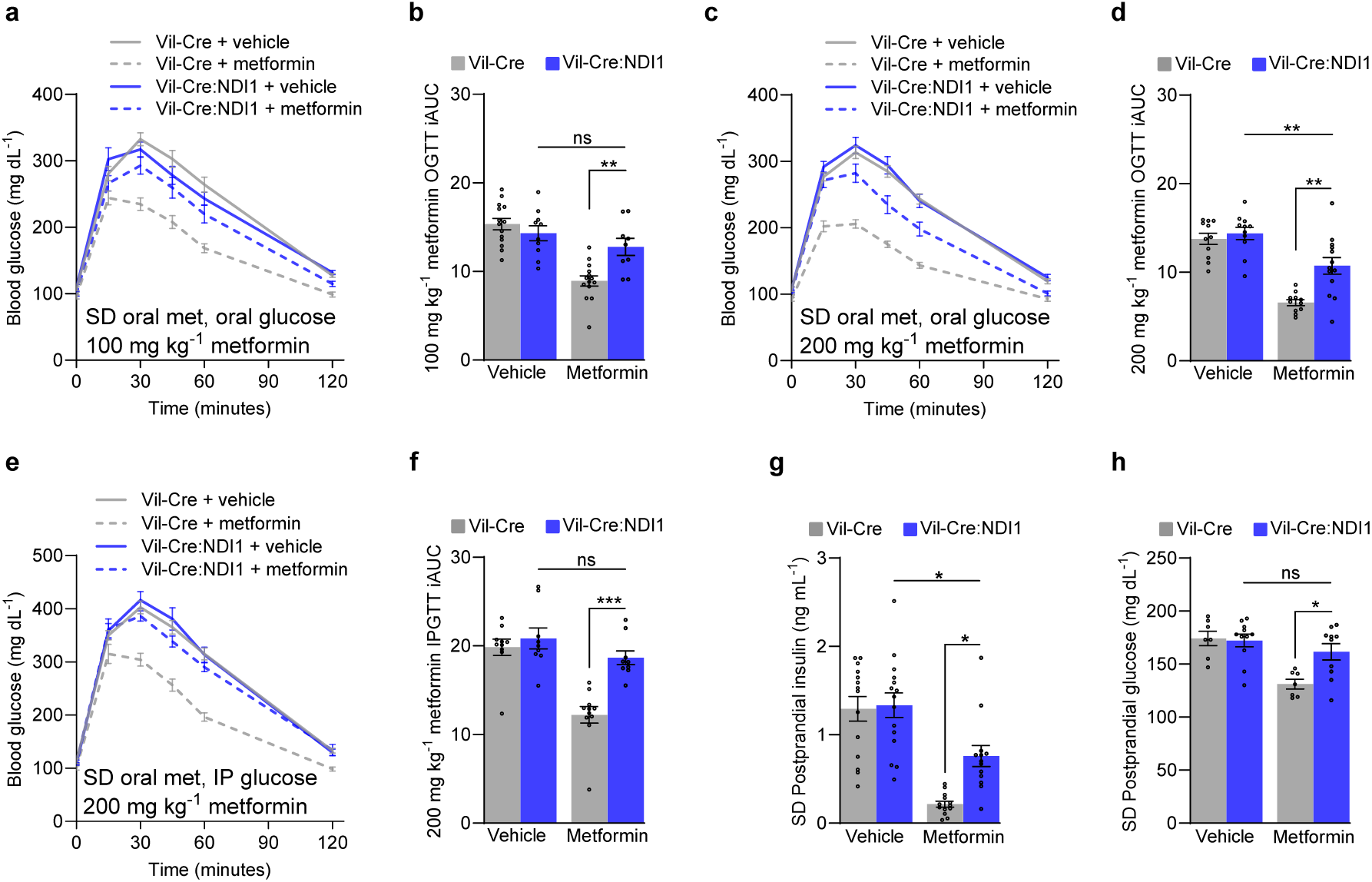
Mitochondrial complex I inhibition in intestinal epithelium is necessary for metformin to improve glycemic control in lean mice. (**a**) Oral glucose tolerance test of standard diet fed mice. Mice were fasted overnight, followed by oral administration of vehicle (water) or metformin (100 mg kg^-1^) and an oral bolus of glucose (2 g kg^-1^) 30 minutes later. (**b**) Incremental area under the curve of (a); Vil-Cre^vehicle^ n=14, Vil-Cre:NDI1^vehicle^ n=10, Vil-Cre^metformin^ n=14, Vil-Cre:NDI1^metformin^ n=9. (**c**) Oral glucose tolerance test of standard diet-fed mice. Vehicle (water) or metformin (200 mg kg^-1^) was orally delivered, followed by an oral bolus of glucose (2 g kg^-1^) 30 minutes later in overnight-fasted mice. (**d**) Incremental area under the curve of (c); Vil-Cre^vehicle^ n=11, Vil-Cre:NDI1^vehicle^ n=11, Vil-Cre^metformin^ n=11, Vil-Cre:NDI1^metformin^ n=13. (**e**) Glucose tolerance test in which standard diet-fed mice were overnight fasted then orally delivered vehicle (water) or metformin (200 mg kg^-1^) then 30 minutes later intraperitoneally injected with glucose (2 g kg^-1^). (**f**) Incremental area under the curve of (e); Vil-Cre^vehicle^ n=10, Vil-Cre:NDI1^vehicle^ n=9, Vil-Cre^metformin^ n=11, Vil-Cre:NDI1^metformin^ n=9. (**g**) Postprandial insulin levels after a fasting-refeeding assay in standard diet-fed mice; Vil-Cre^vehicle^ n=14, Vil-Cre:NDI1^vehicle^ n=15, Vil-Cre^metformin^ n=13, Vil-Cre:NDI1^metformin^ n=13. (**h**) Postprandial glucose levels after a fasting-refeeding assay in standard diet-fed mice; Vil-Cre^vehicle^ n=7, Vil-Cre:NDI1^vehicle^ n=11, Vil-Cre^metformin^ n=7, Vil-Cre:NDI1^metformin^ n=10. SD = standard diet; OGTT = oral glucose tolerance test; IPGTT = intraperitoneal glucose tolerance test; iAUC = incremental area under the curve (arbitrary units). All mice were male and 7-10 weeks of age. In (g) and (h), mice were fasted overnight, followed by an oral dose of vehicle (water) or metformin (200 mg kg^-1^); 30 minutes later, mice were refed ad libitum for 30 minutes, followed by blood collection. Data are presented as mean ± SEM. Statistical significance was determined by Two-way ANOVA with Bonferroni’s correction for multiple comparisons. *P<0.05, **P<0.01, ***P<0.001.

**Figure 4.**
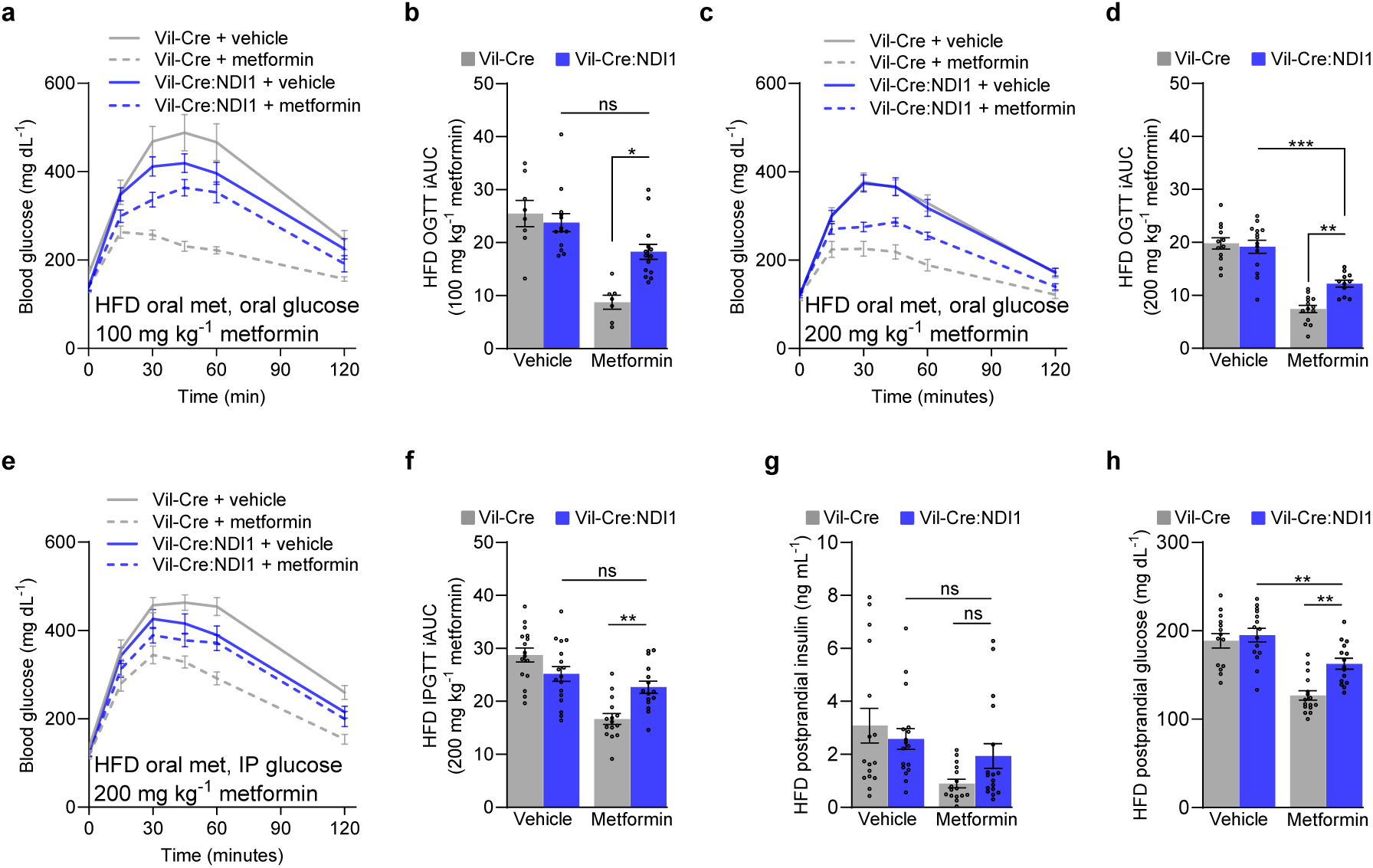
Mitochondrial complex I inhibition in intestinal epithelium is necessary for metformin to improve glycemic control in obese mice. (**a**) Oral glucose tolerance test of diet-induced obese mice fed a high-fat diet for 8-10 weeks. Mice were fasted overnight, followed by oral administration of vehicle (water) or metformin (100 mg kg^-1^) and an oral bolus of glucose (2 g kg^-1^) 30 minutes later. (**b**) Incremental area under the curve of (a); Vil-Cre^vehicle^ n=8, Vil-Cre:NDI1^vehicle^ n=13, Vil-Cre^metformin^ n=7, Vil-Cre:NDI1^metformin^ n=14. (**c**) Oral glucose tolerance test of diet-induced obese mice fed a high-fat diet for 8-10 weeks. Vehicle (water) or metformin (200 mg kg^-1^) was orally delivered, followed by an oral bolus of glucose (2 g kg^-1^) 30 minutes later in overnight-fasted mice. (**d**) Incremental area under the curve of (c); Vil-Cre^vehicle^ n=12, Vil-Cre:NDI1^vehicle^ n=14, Vil-Cre^metformin^ n=14, Vil-Cre:NDI1^metformin^ n=11. (**e**) Glucose tolerance test in which diet-induced obese mice were overnight fasted then orally delivered vehicle (water) or metformin (200 mg kg^-1^) then 30 minutes later intraperitoneally injected with glucose (2 g kg^-1^). (**f**) Incremental area under the curve of (e); Vil-Cre^vehicle^ n=16, Vil-Cre:NDI1^vehicle^ n=17, Vil-Cre^metformin^ n=15, Vil-Cre:NDI1^metformin^ n=15. (**g**) Postprandial insulin levels after a fasting-refeeding assay in diet-induced obese mice; Vil-Cre^vehicle^ n=16, Vil-Cre:NDI1^vehicle^ n=17, Vil-Cre^metformin^ n=16, Vil-Cre:NDI1^metformin^ n=17. (**h**) Postprandial glucose levels after a fasting-refeeding assay in diet-induced obese mice; Vil-Cre^vehicle^ n=14, Vil-Cre:NDI1^vehicle^ n=15, Vil-Cre^metformin^ n=16, Vil-Cre:NDI1^metformin^ n=15. HFD = high-fat diet (60% lard); OGTT = oral glucose tolerance test; IPGTT = intraperitoneal glucose tolerance test; iAUC = incremental area under the curve (arbitrary units). All mice were started on HFD at 8 weeks of age, and experiments were performed after 8-10 weeks of HFD feeding. In (g) and (h), mice were fasted overnight, followed by an oral dose of vehicle (water) or metformin (200 mg kg^-1^); 30 minutes later, mice were refed ad libitum for 30 minutes, followed by blood collection. Data are presented as mean ± SEM. Statistical significance was determined by Two-way ANOVA with Bonferroni’s correction for multiple comparisons. *P<0.05, **P<0.01, ***P<0.001.

Given these findings with acute oral administration of metformin, we next sought to determine whether the attenuation of metformin’s glucose-lowering effect by intestinal NDI1 expression persists under chronic treatment conditions. To this end, diet-induced obese mice were given metformin in their drinking water for two weeks at a dose (1 mg mL⁻¹) recently employed to link PEN2 to metformin’s mechanism of action^75^. Using this dosing regimen, metformin did not lower fasting glucose or improve glucose tolerance in control or Vil-Cre:NDI1 mice (**Extended Data Fig. 7a-c**), indicating that administering metformin in drinking water at 1 mg mL^-1^ is not sufficient to promote glycemic control. This is likely due to the lower and more fluctuating metformin exposure of this method compared to acute oral gavage, which better mimics the pharmacokinetics observed in patients^7^.

Having established the necessity of intestinal mitochondrial complex I inhibition for acutely administered metformin to improve glucose tolerance, we next turned to a related clinical metric of glycemic control: postprandial glucose excursions. Postprandial hyperglycemia is a key feature of type II diabetes and a better predictor of cardiovascular disease and all-cause mortality than fasting glucose^34,35^. Since metformin is known to suppress these meal-induced blood glucose spikes, we sought to determine whether mitochondrial complex I inhibition in intestinal epithelium is necessary for this clinical effect of metformin. To do so, we devised a simple fasting-refeeding assay in which mice were fasted overnight, treated with vehicle or metformin, and refed ad libitum 30 minutes before blood collection. Metformin lowered glucose and insulin levels in refed controls but this effect was attenuated in Vil-Cre:NDI1 animals (**Fig. 3g, h**). Similar effects were seen in diet-induced obese mice, though insulin levels were more variable (**Fig. 4g, h**). Thus, mitochondrial complex I inhibition in intestinal epithelium is necessary for metformin to improve postprandial glucose control.

### Intestine-specific mitochondrial complex I inhibition is essential for metformin to improve pyruvate tolerance

Historically, research on metformin has centered around its effect on hepatic gluconeogenesis, an anabolic pathway responsible for synthesizing glucose from smaller, non-carbohydrate precursors. Pyruvate tolerance tests (PTTs) are commonly used to assess metformin’s impact on this pathway, since pyruvate is a key substrate that the liver converts into glucose through gluconeogenesis^76^. The resulting rise in blood glucose reflects hepatic gluconeogenesis, making the PTT a useful tool for evaluating alterations in this biosynthetic process. However, pyruvate can also be metabolized in the opposite direction to form lactate and regenerate cytosolic NAD^+^ to support glycolysis during electron transport chain impairment^77^. We hypothesized that metformin could improve pyruvate tolerance by enhancing the consumption of pyruvate in the intestines (**Fig. 5a**).

**Figure 5.**
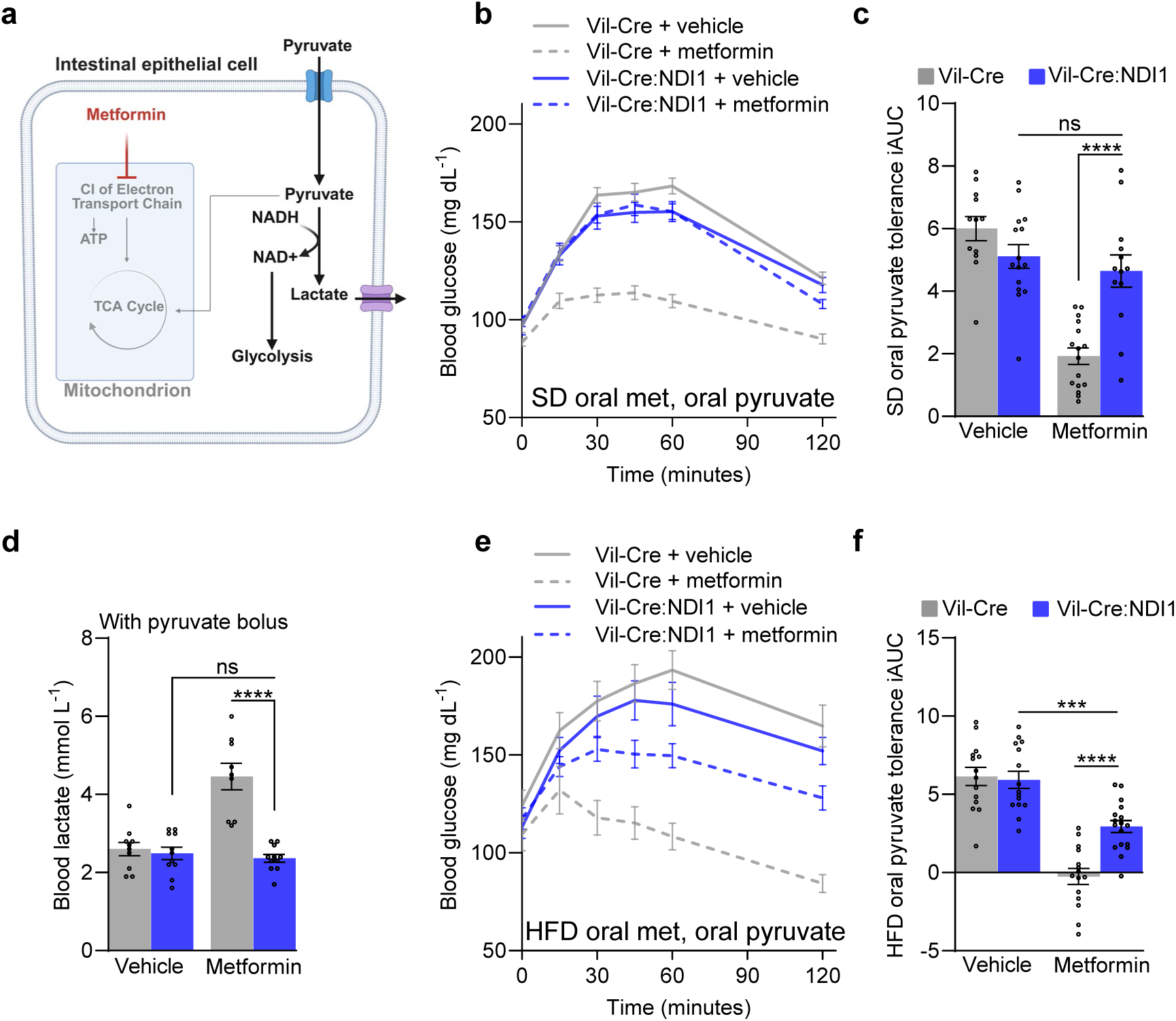
Mitochondrial complex I inhibition in intestinal epithelium is necessary for metformin to improve pyruvate tolerance. (**a**) Model of metformin-induced improvement in pyruvate tolerance. Created with BioRender.com. (**b**) Oral pyruvate tolerance test with metformin treatment on standard diet fed mice. Mice were fasted overnight, followed by oral administration of vehicle (water) or metformin (200 mg kg^-1^) and an oral bolus of pyruvate (2 g kg^-1^) 30 minutes later. (**c**) Incremental area under the curve of (a); Vil-Cre^vehicle^ n=12, Vil-Cre:NDI1^vehicle^ n=15, Vil-Cre^metformin^ n=16, Vil-Cre:NDI1^metformin^ n=13. (**d**) Blood lactate levels in overnight fasted mice on standard diet. Mice were orally administered vehicle (water) or metformin (200 mg kg^-1^), followed by an oral gavage of pyruvate (2 g kg^-1^) 30 minutes later; 30 minutes after pyruvate administration, blood lactate was measured; Vil-Cre^vehicle^ n=10, Vil-Cre:NDI1^vehicle^ n=11, Vil-Cre^metformin^ n=9, Vil-Cre:NDI1^metformin^ n=11. (**e**) Oral pyruvate tolerance test with metformin of high-fat diet fed mice. Mice were fasted overnight, followed by oral administration of vehicle (water) or metformin (200 mg kg^-1^) and an oral bolus of pyruvate (2 g kg^-1^) 30 minutes later. (**f**) Incremental area under the curve of (d); Vil-Cre^vehicle^ n=16, Vil-Cre:NDI1^vehicle^ n=17, Vil-Cre^metformin^ n=15, Vil-Cre:NDI1^metformin^ n=15. SD = standard diet; HFD = high-fat diet (60% lard); iAUC = incremental area under the curve (arbitrary units). All mice were male. For SD-fed mice, pyruvate tolerance tests were performed on 7–10-week-old animals; blood lactate measurements were performed on 9-12-week-old animals. For HFD-fed mice, HFD was started at 8 weeks of age and pyruvate tolerance tests were performed after 8-10 weeks of HFD feeding. Data are presented as mean ± SEM. Statistical significance was determined by Two-way ANOVA with Bonferroni’s correction for multiple comparisons. ***P<0.001, ****P<0.0001.

Accordingly, we found that intestinal NDI1 expression significantly attenuates metformin-induced improvement in pyruvate tolerance in both lean and obese mice (**Fig. 5b, c, e, f; Extended Data Fig. 8a-d**). Consistent with this, exogenous pyruvate strongly enhances metformin-induced lactate production in a manner that requires intestinal mitochondrial complex I inhibition (**Fig. 5d**). Together, these findings show that the improved pyruvate tolerance seen with metformin treatment is not strictly due to a direct effect on the liver. Instead, metformin inhibits mitochondrial complex I in intestinal epithelium to trigger a metabolic shift that redirects exogenous pyruvate away from hepatic gluconeogenesis toward the intestines to support glycolysis.

### Phenformin and berberine inhibit mitochondrial complex I in intestinal epithelium to promote glycemic control

While metformin is currently the only FDA-approved biguanide drug, another biguanide, phenformin (**Fig. 6a**), was formerly used for blood glucose control prior to its withdrawal due to increased risk of lactic acidosis. We assessed whether intestinal mitochondrial complex I inhibition is necessary for phenformin to improve glucose tolerance. As expected, the blood glucose lowering effect of phenformin (100 mg kg^-1^) is attenuated in Vil-Cre:NDI1 mice (**Fig. 6b, c**). Thus, biguanides target mitochondrial complex I in intestinal epithelium to promote glycemic control.

**Figure 6.**
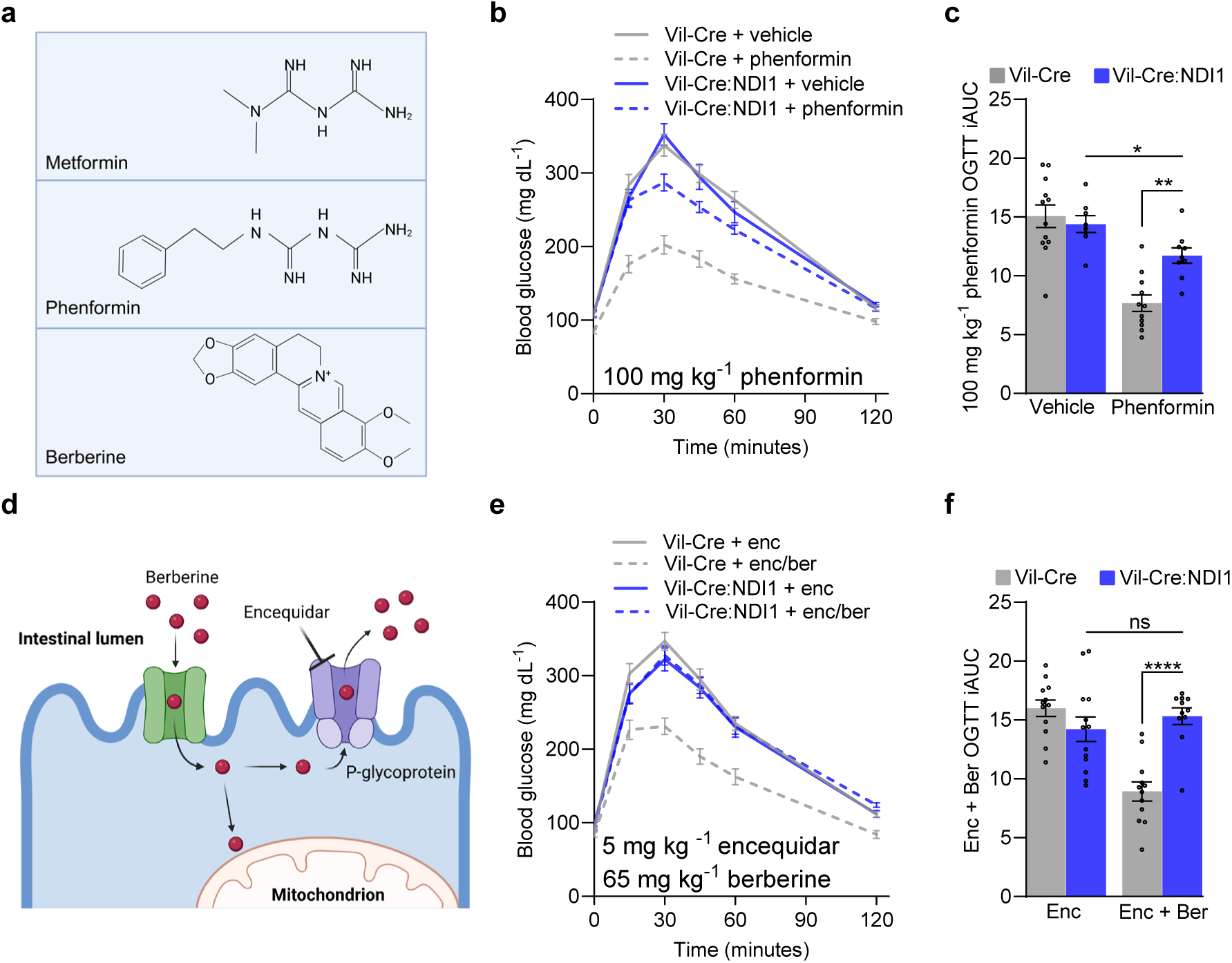
Phenformin and berberine inhibit mitochondrial complex I in intestinal epithelium to improve glucose tolerance. (**a**) Chemical structures of metformin, phenformin, and berberine. Created with BioRender.com. (**b**) Oral glucose tolerance test with phenformin. Mice were fasted overnight, followed by oral administration of vehicle (water) or phenformin (100 mg kg^-1^) and an oral bolus of glucose (2 g kg^-1^) 30 minutes later. (**c**) Incremental area under the curve of (b); Vil-Cre^vehicle^ n=12, Vil-Cre:NDI1^vehicle^ n=8, Vil-Cre^phenformin^ n=11, Vil-Cre:NDI1^phenformin^ n=9. (**d**) Model of berberine’s mechanism of action in the intestinal epithelium. Created with BioRender.com. (**e**) Oral glucose tolerance test with berberine (65 mg kg^-1^) cotreated with encequidar (5mg kg^-1^). Mice were fasted overnight, followed by oral administration of encequidar (5 mg kg^-1^) or encequidar (5 mg kg^-1^) + berberine (65 mg kg^-1^) and an oral bolus of glucose (2 g kg^-1^) 30 minutes later. (**f**) Incremental area under the curve of (e); Vil-Cre^enc^ n=12, Vil-Cre:NDI1^enc^ n=13, Vil-Cre^enc/ber^ n=12, Vil-Cre:NDI1^enc/ber^ n=11. All mice were male and 7-10 (b, c) or 9-12 (e, f) weeks of age. Data in (c) and (f) are presented as mean ± SEM. OGTT = oral glucose tolerance test, iAUC = incremental area under the curve (arbitrary units). Statistical significance was determined by Two-way ANOVA with Bonferroni’s correction for multiple comparisons. **P<0.01, ****P<0.0001.

Given that biguanides originate from guanidine, a natural compound in French Lilac^78^, we wondered whether any other natural compounds selectively target mitochondrial complex I in the intestines. Putative complex I inhibitors found in nature include rotenone, annonacin, and berberine, which are plant-derived allelochemicals with antifeedant and insecticidal activities^79–83^. While rotenone and annonacin are neurotoxic to humans^81,84,85^, berberine is marketed as a dietary supplement to improve metabolic homeostasis. Clinical data is limited but suggests that berberine and metformin have overlapping pharmacological profiles, including gastrointestinal side effects^86^. Berberine, which is structurally unrelated to biguanides (**Fig. 6a**), is a potent mitochondrial complex I inhibitor in vitro^87^. However, its extremely low oral absorption (due to P-glycoprotein-mediated efflux into the intestinal lumen)^87,88^, has made it challenging to ascertain a mechanism of action in vivo. Given berberine’s gut-restricted biodistribution, we hypothesized that it preferentially targets mitochondrial complex I in the intestinal epithelium to promote glycemic control.

We found that a single oral dose of berberine (1000 mg kg^-1^) improves glucose tolerance in controls but not Vil-Cre:NDI1 mice (**Extended Data Fig. 9a, b**), indicating intestine-selective mitochondrial complex I inhibition is necessary for berberine to lower blood glucose levels. By co-treating mice with encequidar, an intestine-specific p-glycoprotein inhibitor (**Fig. 6d**)^89^, we observed a far more dramatic antihyperglycemic effect of berberine (65 mg kg^-1^) in control animals (**Fig. 6e, f**). Whereas, in contrast to biguanides, the glucose-lowering effect of berberine was completely blocked by intestinal NDI1 expression. These results indicate that intestine-specific mitochondrial complex I inhibition is the principal mechanism by which berberine acutely lowers blood glucose and further underscores the therapeutic utility of targeting complex I in the gut.

## Discussion

In this study, we show how metformin exerts multiple clinical effects through selective inhibition of mitochondrial complex I in the intestinal epithelium. This mechanism suppresses citrulline synthesis and drives increased glucose uptake, glycolysis, and lactate production in the intestines, leading to improved glucose tolerance, pyruvate tolerance, and postprandial glycemia, along with increases in Lac-Phe and GDF15, biomarkers of mitochondrial stress that have been linked to metformin-associated body weight regulation. While intestinal complex I inhibition is necessary for these effects, the incomplete resistance to glucose lowering observed in both intestinal NDI1 mice in this study and whole-body NDI1 mice in our previous work^57^ suggests that metformin also acts through additional targets in other organs, a possibility supported by existing literature^2,7,90^. Alternatively, NDI1 may not fully compensate for the suppression of mammalian complex I due to its inability to pump protons or generate superoxide. Indeed, NDI1 confers modestly greater resistance when metformin is administered at low doses (**Fig. 3a, b**; **Fig. 4a, b**). In either case, our results establish that inhibition of mitochondrial complex I in intestinal epithelium is therapeutically indispensable for not only metformin, but also phenformin and the structurally unrelated molecule berberine. This shared mechanism highlights the intestines as a major site of action for these therapeutics, underscoring the clinical utility of gut-selective mitochondrial complex I inhibition in promoting glycemic control.

Pertaining to berberine, its affinity for P-glycoproteins is consistent with its biological function as a mitochondrial toxin that defends against herbivores. P-glycoproteins are promiscuous efflux pumps that extrude potentially harmful xenobiotics into the intestinal lumen^91^. We speculate that the intestine-restricted activity of berberine is necessary for both its safety and efficacy. By limiting systemic absorption, P-glycoproteins may enable berberine to reach therapeutic levels in the gut while reducing the risk of toxicity in other tissues—a property likely to be important given that berberine is a much stronger inhibitor of mitochondrial complex I (IC_50_ ≈ 15 μM) compared to phenformin (IC_50_ ≈ 430 μM) and metformin (IC_50_ ≈ 19,400 μM)^53,87^.

Intestine-specific inhibition of mitochondrial complex I also accounts for metformin-induced citrulline depletion. A key question is whether the observed citrulline depletion in patients taking metformin impacts metabolic health outcomes. Circulating citrulline is the primary precursor to nitric oxide, a potent vasodilator essential for muscle perfusion during exercise^48^. Citrulline is upregulated by exercise in humans^92^ and is the dominant ingredient in many pre-workout supplements because it enhances muscle perfusion and exercise performance^93^. In contrast, metformin impairs muscle hypertrophy and aerobic capacity in response to exercise^94,95^. A direct inhibitory effect of metformin on muscle mitochondria is unlikely due to the low concentration of metformin in this tissue (**Fig. 1e**)^12,52^. However, it is plausible that metformin-induced citrulline depletion, and thus reduced nitric oxide production, underlies the blunted exercise benefits caused by metformin. Indeed, supplementing with citrulline could be a straightforward and scalable solution to support exercise adaptation in patients taking metformin. Should this approach be effective, it may be possible to more safely investigate the geroprotective potential of metformin^96^, enabling the rigorous clinical evaluation of its ability to delay multiple age-related diseases while mitigating harmful side effects.

Much of the literature has focused on plasma concentrations of metformin when evaluating therapeutically relevant exposures in preclinical models^10,97^. However, such an emphasis overlooks key aspects of metformin pharmacology, as plasma levels are an unreliable indicator of the drug’s distribution and accumulation in tissues^12,52^. Indeed, many rodent studies have delivered metformin via portal vein infusions and intraperitoneal injections, which bypass the gut, or have administered metformin in drinking water. While the latter approach more closely resembles clinical practice, it still exposes animals to a variable, low-level of metformin rather than the characteristic “peak-and-trough” exposure of patients^7,98,99^. One particularly important consideration is that orally delivered metformin exhibits “flip-flop” pharmacokinetics, in which the rate of its appearance in plasma is slower than its rate of elimination^98,100,101^. This feature has major implications for understanding what constitutes a tissue-specific therapeutic dose of metformin.

The “flip-flop” pharmacokinetic profile of metformin arises from the localization of its transporters in the intestinal epithelium. When administered orally, metformin is rapidly taken up from the intestinal lumen into enterocytes via several apical membrane transporters (PMAT, OCTN1, OCT3, THTR-2, and SERT). However, its export from enterocytes into the bloodstream is constrained by a single basolateral transporter, OCT1^102^. As a result of this bottleneck, metformin concentrations in the intestinal epithelium can exceed plasma levels by up to 300-fold and are estimated to be 10 to 100 times higher than in the liver, reaching concentrations well into the millimolar range^11–13^.

Accordingly, while it is generally accepted that metformin reduces endogenous glucose production through direct inhibition of hepatic gluconeogenesis, clinical evidence shows that endogenous glucose production is not suppressed by metformin in patients with mild hyperglycemia or early-stage type II diabetes^17–20^, who represent the vast majority of metformin users^103^. Metformin has even been found to elevate endogenous glucose production among these patient groups^18,19^. Instead, glycemic control is achieved by increasing the rate of glucose clearance^18^. Importantly, this increase in glucose clearance is observed across diverse patient populations, ranging from normoglycemic subjects^17,18,21,104^ to those with overt type II diabetes^25,105–108^. Therefore, enhanced glucose clearance is a robust, clinically reproducible effect of metformin.

Our findings show that metformin enhances glucose clearance by inhibiting mitochondrial complex I in the intestinal epithelium. This inhibition co-opts the intestines to function as a glucose sink, drawing in excess glucose and channeling it into glycolysis. While this mechanism is central to metformin’s therapeutic effect, important questions remain. Our data and the work of others indicate that, in addition to intestinal complex I, metformin engages other targets, and its glucoregulatory effects vary with route of administration, tissue distribution, and the stage of type II diabetes^2,5,17,90^. For example, the resistance conferred by NDI1 is more pronounced at lower metformin doses and when glucose is administered intraperitoneally (**Fig. 3a-f**, **Fig. 4a-f**), suggesting metformin may have distinct effects on glucose in the intestinal lumen. The gut microbiota may be particularly relevant given their direct exposure to metformin and sensitivity to host metabolism^109^. Elucidating metformin’s extra-intestinal mechanisms will also be critical for fully defining its therapeutic profile and optimizing clinical use.

## Methods

### Animals

Villin-Cre mice (B6.Cg-Tg(Vil1-cre)1000Gum/J) were obtained from the Jackson Laboratory (Strain #: 021504). NDI1^LSL^ mice have been described previously. Homozygous Vil-Cre mice were crossed with heterozygous NDI1^LSL^ mice to generate Vil-Cre control mice and Vil-Cre:NDI1 mice. The study used littermate male Vil-Cre and Vil-Cre:NDI1 mice. Mice were housed in the Northwestern Center for Comparative Medicine vivarium in a temperature- and humidity-controlled room (23^°^C with 30-70% humidity range) with a 12-hour light/dark cycle. Mice were monitored by research staff, as well as Northwestern Comparative Medicine animal care technicians and veterinary staff. Mice were group-housed with free access to water and standard chow (Envigo/Teklad LM-485) or a 60% lard high-fat diet (Research diets, D12492i). Where noted in the main text and figure legends, overnight fasts were 16 to 18 hours with free access to water. After relevant procedures, mice were refed immediately. For all procedures performed on standard chow diet (SD) fed animals, mice were 7 to 12 weeks of age. For animals fed a high-fat diet (HFD), the HFD switch was performed at 8 weeks of age, and experiments were performed after 8 to 12 weeks of HFD feeding. Metformin, phenformin, berberine, and encequidar were administered via 18G x 50mm curved oral gavage needles (GavageNeedle, Cat. #AFN1850C) attached to BD syringes with Luer-Lok Tips (Thermo Fisher Scientific, Cat. #14-823-30). The Northwestern University Institutional Animal Care and Use Committee (IACUC) reviewed and approved all animal procedures used in this study.

### Preparation of biguanides, berberine and encequidar

Metformin tablets (Metformin Hydrochloride, Granules Pharmaceuticals Inc., NDC 70010-064-01) were crushed to powder with a mortar and pestle and dissolved in water. Phenformin powder (Cayman Chemical, Item #14997) and berberine powder (Cayman Chemical, Item # 10006427) were dissolved directly into water. Encequidar (Cayman Chemical, HM30181, Item # 32873) was dissolved in 0.5% DMSO. For encequidar/berberine co-treatment, both drugs were dissolved in 0.5% DMSO.

### Tissue quantification of metformin

Five male C57BL/6J mice (10 weeks old) were fasted overnight (16-18 hours), followed by an oral gavage of metformin (200 mg kg^-1^). One hour later, blood was collected via tail vein nick into microhematocrit tubes (Thermo Fisher Scientific, no. 22-362-566), and mice were euthanized. The intestine (jejunum), muscle (soleus), and liver tissue were harvested and frozen on dry ice. Blood was centrifuged in 1.5 ml microcentrifuge tubes at 10,000 x g at 4°C for 10 minutes to collect plasma. Plasma and tissue were stored at - 80°C until further processing. Tissue was homogenized using a QIAGEN TissueRuptor II in cold acetonitrile/water (80/20, v/v) with a 2 uM metformin-D6 (Cayman Chemical, no. 16921) internal standard (80 ul acetonitrile/water per 1 mg tissue). 10 ul of plasma was added to 90 ul acetonitrile/water (80/20, v/v). The samples underwent three freeze/thaw cycles, then were centrifuged at 17,000 x g at 4°C for 10 minutes. Supernatants containing metformin and soluble metabolites were collected.

As previously^57^, a standard curve was made using metformin concentrations ranging from 25 nM to 200 uM (Cayman Chemical, no. 13118) in 80/20 acetonitrile/water spiked with 2 uM metformin-D6 (Cayman Chemical, no. 16921). High-performance liquid chromatography and triple quadrupole tandem mass spectrometry (HPLC-MS/MS) analyzed the standards and samples. The system consists of a TSQ (Thermo Fisher Scientific) in line with an electrospray ion source (ESI) and Vanquish (Thermo Fisher Scientific) UHPLC with a binary pump, degasser, and auto-sampler outfitted with an XBridge C18 column (Waters, dimensions of 2.1 mm by 50 mm, 3.5 uM). Solute separation was achieved through isocratic elution with the mobile phase containing 0.1% formic acid in acetonitrile/water (65/35, v/v) at 0.15 ml/min. The capillary ESI was set to 300°C in positive mode, with sheath gas at 35 arbitrary units, auxiliary gas at five arbitrary units, and the spray voltage at 3.5 kV. Selective reaction monitoring of the protonated precursor ion and the related product ions for metformin and metformin-D6 [mass/charge ratio (m/z) 130.15 → 71, 136.15 → 77, respectively] was performed. The standard curve was calculated from the peak area ratio of targets to internal standard with a linear regression R^2^ = 0.99998. Data were acquired using Xcalibur 4.1 software and analyzed via TraceFinder 4.1 (Thermo Fisher Scientific). Tissue and plasma metformin concentrations were calculated using the standard curve after normalizing to the internal metformin-D6 standard.

### Glucose and pyruvate tolerance tests and blood lactate quantification

For glucose and pyruvate tolerance tests, mice were fasted overnight (16-18 hours) followed by an oral gavage of vehicle or drug (i.e., metformin, phenformin, berberine, or encequidar + berberine). Thirty minutes later, fasting blood glucose was measured via a tail vein nick with a Contour Next glucose test meter. Immediately after the blood glucose measurement, mice were given an oral gavage or intraperitoneal (IP) injection of glucose or pyruvate (2 g kg^-1^). For oral dosing, glucose/pyruvate was dissolved in water; for IP dosing, glucose/pyruvate was dissolved in sterile PBS. Blood glucose was measured 15, 30, 45, 60 and 120 minutes following glucose/pyruvate administration. Glucose tolerance tests were performed the same way after two weeks of metformin treatment in drinking water, except that mice did not receive an acute dose of metformin on the test day. The incremental area under the curve was calculated using GraphPad Prism software v10.4.1.

Similarly, for blood lactate measurements, mice were fasted overnight (16-18 hours) followed by oral gavage of vehicle or drug (i.e., metformin or phenformin). Thirty minutes later, mice were given an oral bolus of water or glucose/pyruvate (2 g kg^-1^). After an additional thirty minutes, blood lactate was determined via a tail vein nick with a Nova Biomedical Lactate Plus meter.

### Body weight and food intake measurements

Mice were placed on a high-fat diet beginning at 8 weeks of age. After 4 weeks on the diet, they were single-housed for 3 days before undergoing a 4-day oral gavage conditioning period, during which they received one daily gavage of water. Following conditioning, mice and their food intake were measured daily for two weeks. During this monitoring period, mice received a daily oral gavage of either vehicle (water) or metformin (200 mg/kg).

### Fasting-refeeding assay

Mice were fasted overnight (16-18 hours), followed by single-housing and an oral gavage of vehicle (water) or metformin (200 mg kg^-1^). Thirty minutes later, mice were refed ad libitum for half an hour. After refeeding, blood glucose was measured via a tail vein nick with a Contour Next blood glucose meter. For insulin measurement, additional blood was collected into microhematocrit tubes (Thermo Fisher Scientific, no. 22-362-566). Following blood collection, mice were returned to their original cages. Blood within microhematocrit tubes was transferred to 1.5 ml microcentrifuge tubes on ice and then spun down at 10,000 x g at 4°C for 10 minutes. Plasma was collected and stored at -80^°^C until insulin was quantified using the Ultra-Sensitive Mouse Insulin ELISA kit (Crystal Chem, no. 90080) according to the manufacturer’s instructions.

### Plasma GDF15 quantification

Mice were fasted overnight (16-18 hours), followed by an oral gavage of vehicle (water) or metformin (200 mg kg^-1^). Eight hours later, blood was collected, and plasma was acquired/stored as in the fasting-refeeding assay above. GDF15 was measured using the Mouse/Rat GDF15 Quantikine ELISA kit (cat. # MDG150) according to the manufacturer’s instructions.

### Serum quantification of Lac-Phe and citrulline

Mice were fasted overnight (16-18 hours), followed by an oral gavage of vehicle (water) or metformin (200 mg kg^-1^). Thirty minutes later, mice were given an oral glucose bolus (2 g kg^-1^). Blood was collected two hours after the glucose bolus, and plasma was acquired/stored as above. 5 µL of plasma was added to 170 µL of cold 100% HPLC-grade methanol to extract metabolites. Samples were vortexed and then incubated on dry ice for 5 minutes. Samples were then centrifuged for 10 minutes at 16,000 x g at 4°C. Following centrifugation, 50 µL of supernatant was added to 50 µL of cold methanol/water (80/20, v/v) with a thymine-D4 internal standard at 200ng/mL. Samples were then vortexed and centrifuged for 10 minutes at 16,000 x g at 4°C. 100 ul of supernatant was transferred to HPLC tubes for metabolite quantification as previously reported ^110^. The supernatant was collected for LC–MS analysis. LC was performed on an Xbridge BEH amide HILIC column (Waters) with a Ultimate3000 HPLC system (ThermoFisher). Solvent A was 95:5 water: acetonitrile with 20 mM ammonium acetate and 20 mM ammonium hydroxide at pH 9.4. Solvent B was acetonitrile. The gradient used for metabolite separation was 0 min, 90% B; 2 min, 90% B; 3 min, 75%; 7 min, 75% B; 8 min, 70% B, 9 min, 70% B; 10 min, 50% B; 12 min, 50% B; 13 min, 25% B; 14 min, 25% B; 16 min, 0% B, 21 min, 0% B; 21 min, 90% B; and 25 min, 90% B. MS analysis was performed on an expoloris240 mass spectrometer (ThermoFisher) in polarity switching mode, scanning an *m*/*z* range of 70 to 1,000. Data were analyzed using El-MAVEN Software (Elucidata; elucidata.io)^111^.

### Jejunum metabolomics & plasma U-^13^C_6_-glucose tracing

Mice were fasted overnight (16-18 hours), followed by an oral gavage of vehicle (water) or metformin (200 mg kg^-1^). One hour later, mice were euthanized with isoflurane, and jejunum was harvested. Before freezing on dry ice, luminal contents were removed by flushing with PBS and applying gentle pressure. Samples were stored at -80°C until further processing. Jejunum was mechanically homogenized in 20 ul cold acetonitrile/water (80/20, v/v) per 1 mg tissue and freeze/thawed three times before spinning down at 10,000 x g for 10 minutes at 4°C. Supernatants containing soluble metabolites were collected, and HPLC-MS/MS was performed. Metabolite separation was achieved through gradient elution, and data were acquired using Xcaliber software (v4.1 Thermo Fisher Scientific). Data analysis was performed using MetaboAnalyst v5.0^112^. Metabolite abundance was normalized to the total ion count for each sample. Hierarchical clustering (Ward) was performed with the top 50 differentially abundant metabolites (ANOVA) and shown as a heatmap.

For U-^13^C_6_-glucose tracing, overnight fasted mice were given an oral gavage of vehicle (water) or metformin (200 mg kg^-1^) followed by an oral dose of U-^13^C_6_-glucose (2 g kg^-1^) thirty minutes later. Thirty minutes after the U-^13^C_6_-glucose administration, blood was collected via a tail vein nick and plasma acquired/stored as above. Plasma was diluted 1:100 (v/v) in cold acetonitrile/water (80/20, v/v) and freeze/thawed three times before centrifugation at 10,000 x g for 10 minutes at 4°C. Supernatants were collected and HPLC-MS/MS was performed. The m+3 lactate / m+6 glucose ratio was calculated on a per sample basis with ion counts.

### Human citrulline and metabolomic analysis

Publicly available human metabolomics data were obtained from Rotroff et al.^41^ and Aleidi et al.^42^, both released under a CC-BY license. For Rotroff et al., relative citrulline levels from time point A (before metformin) and time point C (after metformin) were obtained from supplementary data sheet 4 of the original manuscript. All values were divided by 100 prior to visualization to scale the y-axis. The volcano plot was generated using metabolite levels from supplementary data sheet 4 with a custom R script (available on Zenodo); all unknown metabolites were removed before analysis. Glucose tolerance data from Rotroff et al. was obtained from supplementary data sheet 3 of the original manuscript. Relative citrulline levels from Aleidi et al. were obtained from supplementary data sheet 3 of the original manuscript. Notably, the original data from Aleidi et al. are presented in technical duplicate. We averaged these values and present only biological replicates.

### RNA sequencing

Mice were fasted overnight (16-18 hours), followed by an oral gavage of vehicle (water). One hour later, they were euthanized with isoflurane, and the jejunum was harvested. Luminal contents were removed by flushing with PBS and applying gentle pressure, and jejunal tissue was flash-frozen on dry ice. Tissue samples were stored at -80^°^C until RNA was extracted according to the Zymo Research Direct-zol^TM^ RNA MiniPrep instructions with TriReagent kit (Cat. #R2051-A).

RNA sequencing was performed as previously^57^. Briefly, RNA was quantified, and its quality was determined using an Agilent 4200 TapeStation with the RNA ScreenTape System (Agilent Technologies). RNA libraries were prepared using the NEBNext Ultra DNA Library Prep Kit for Illumina (NEB E7370L). Library quality was assessed using the TapeStation 4200 High Sensitivity DNA tapes (Agilent Technologies). Dual-indexed libraries were pooled, and single-end sequenced with an Illumina NextSeq2000 instrument for 100 cycles. BCL Convert v1.2.0 was used to generate FASTQ files. Samples were processed via the publicly available nf-core/rnaseq pipeline v3.12.0 implemented in Nextflow v23.04.3 using Singularity v3.8.1. Minimal command: nextflow run nf-core/rnaseq - r ‘3.12.0’ - profile nu_genomics -- genome ‘GRCm38’ -- additional_fasta ‘S288C_YML120C_NDI1_genomic.fasta’ -- star_index false. Reads were trimmed with trimGalore! v0.6.7 and aligned to the hybrid genome incorporating the NDI1 sequence with STAR v2.6.1d. Gene-level assignments were made with salmon v1.10.1.

Data analysis was performed using custom scripts in R v4.4.0 with DESeq2 v1.46.0. A local gene dispersion model and Wald tests were used for pairwise comparisons. An α threshold of 0.05 was applied for differential expression analysis.

### [18F]FDG-PET/CT

Mice were fasted overnight (16-18 hours), followed by an oral gavage of vehicle (water) or metformin (200 mg kg^-1^). Thirty minutes later, mice were injected via the tail vein with 18F-fluorodeoxyglucose [18F]FDG (∼10.5 MBq) (Sofie Biosciences Inc, Dulles, VA). Following 40 minutes of awake incubation, mice were anesthetized with 1-3% isoflurane and imaged by positron emission tomography (PET) and computed tomography (CT) on the NanoScan8 PET/CT system (Mediso, Hungary). Images were analyzed using ITK-SNAP (3) software. Standard uptake values (SUVs) were internally normalized to the brain, and intestinal FDG accumulation was determined by calculating intestinal volume with SUV>0.45.

### 2-deoxyglucose uptake assay

Mice were fasted overnight (16-18 hours), followed by an oral gavage of vehicle (water) or metformin (200 mg kg^-1^). Thirty minutes later, mice were intraperitoneally injected with glucose (2 g kg^-1^) and 2-deoxyglucose (50 mg kg^-1^). One hour later, mice were euthanized, and the jejunum was harvested. Before freezing on dry ice, luminal contents were removed by flushing with PBS and applying gentle pressure. Samples were stored at -80°C until further processing. The following procedure was adapted from the Promega Glucose Uptake-Glo kit (Cat. #J1342). Jejunum was mechanically homogenized in 10 uL Stop Buffer per mg tissue. Following homogenization, an equal volume of Neutralization Buffer was added and the samples were vortexed. Samples were then freeze/thawed once and centrifuged for 5 minutes at 10,000G to pellet cell debris. Supernatant was transferred to a fresh Eppendorf tube and diluted 1:10 in PBS. Relative 2-deoxyglucose-6-phosphate levels were determined by following the manufacturer’s instructions.

### Statistical Analysis

Except for metabolomics and RNA-seq, data analysis and visualization were performed in GraphPad Prism v10.4.1. All data points are biological replicates. The statistical methods for each experiment are described in the figure legends.

## Data Availability

The manuscript contains all the data required to evaluate its conclusions. Transcriptomic data has been deposited to GEO (accession: GSE293164). Metabolomic data has been deposited in the Metabolomics Workbench (Study ID: ST003841). Code used for human metabolomic and mouse RNA-sequencing analyses are available on Zenodo (DOI: https://10.5281/zenodo.15122149) and will be made publicly accessible upon formal publication.

## Acknowledgements

The authors thank several core facilities at Northwestern—Feinberg School of Medicine, including the Pulmonary NextGen Sequencing Core and the Robert H Lurie. Comprehensive Cancer Center Metabolomics Core. They also thank Daniele Procissi and Lara Leoni, who performed the MicroPET/CT imaging at the Northwestern University Center for Translational Imaging, Small Animal Molecular Imaging (RRID:SCR_017878) Core, which was supported by NCI CCSG P30 CA060553 awarded to the Robert H. Lurie Comprehensive Cancer Center. We thank Anas Abdel Rahman of Alfaisal University and Shereen Aleidi of the University of Jordan for helpful correspondence about their human metabolomics data. We also thank the staff of the Center of Comparative Medicine at Northwestern University.

## Author Contributions

Z.L.S. and N.S.C. conceived of the study, designed experiments, interpreted data, and wrote the manuscript. Z.L.S. conducted most experiments and performed data analysis. C.R.R. created the NDI1 mouse model, contributed to experiments, and provided scientific expertise. R.P.C., A.R.K. and K.B.D. contributed to experiments. R.A.G. created the RNA-seq pipeline and aided with computational analysis. J.L.E.B. and S.M.D. generated a portion of the metabolomic data and aided with analysis.

## Materials & Correspondence

Correspondence regarding mouse lines can be made to Navdeep Chandel.

## Extended Data Figures and Legends

**Extended Data Figure 1.**
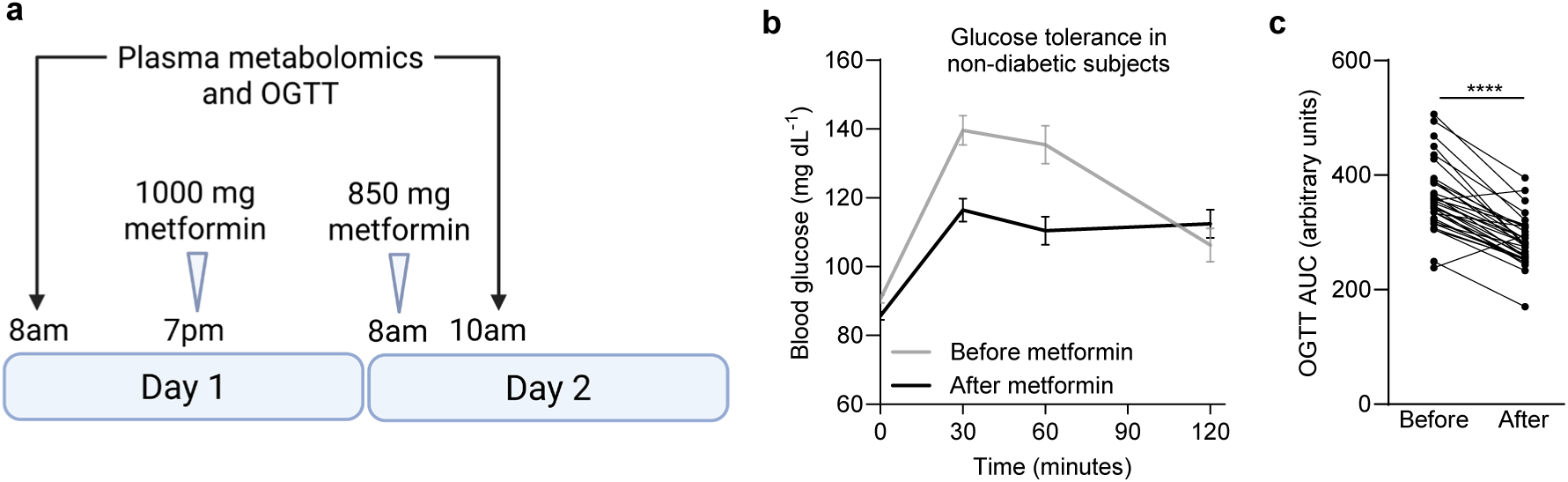
Metformin acutely improves glucose tolerance in non-diabetic humans. (**a**) Schematic illustrating sample collection and metformin dosing in the Rotroff et al. cohort (same as Figure 1a); related to panels (b) and (c). Created with BioRender.com. (**b**) Glucose tolerance test in the Rotroff et al. cohort of non-diabetic subjects before and after metformin treatment. (**c**) Area under the curve of (b); n=33. OGTT = oral glucose tolerance test. Statistical significance for (c) was determined by Paired t test. ****P<0.0001.

**Extended Data Figure 2.**
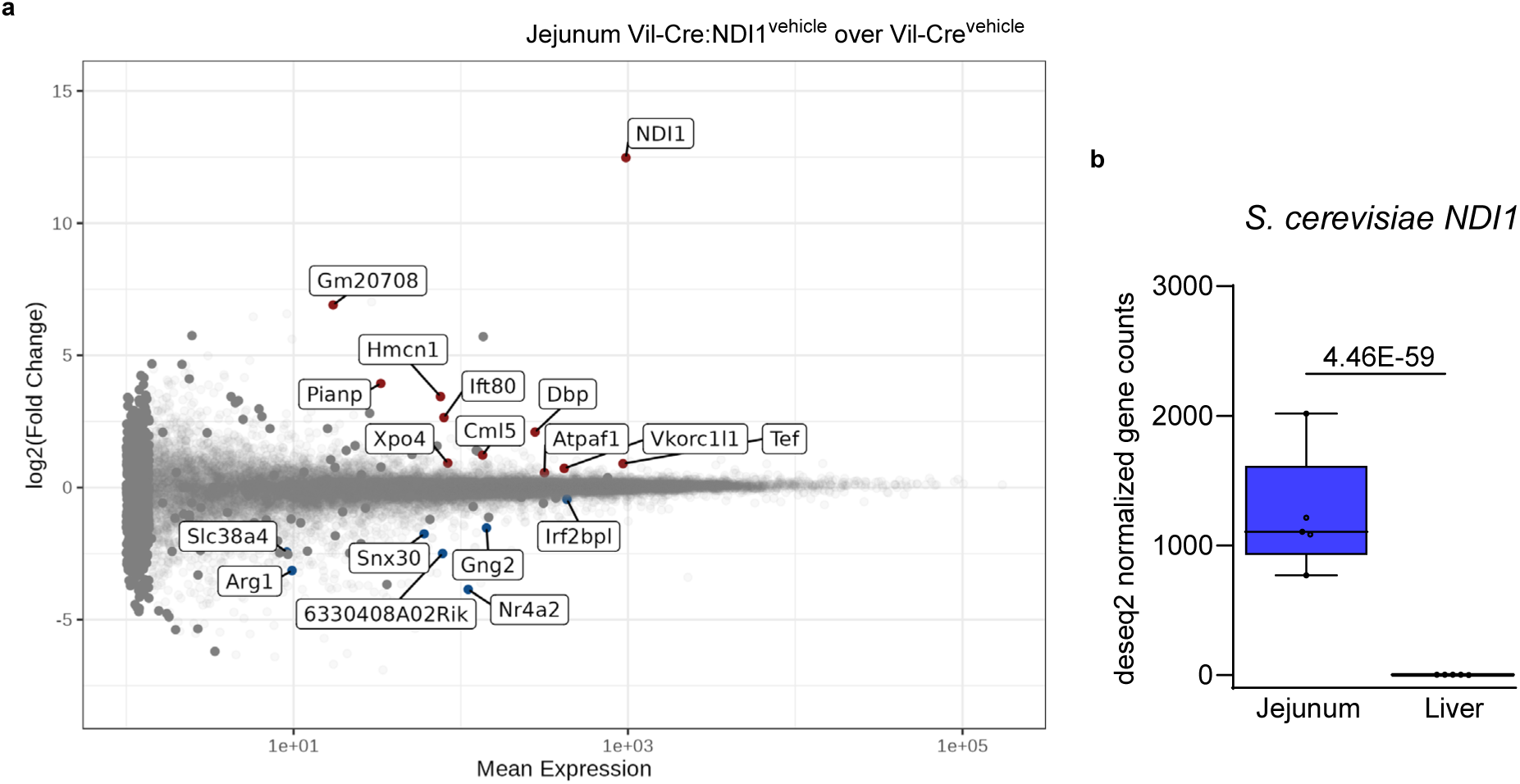
NDI1 has minimal effect on the intestinal transcriptome and is not expressed in liver of Vil-Cre:NDI1 mice. (**a**) MA plot showing differential gene expression in the jejunum of overnight fasted mice one hour after oral gavage of vehicle; Vil-Cre n=5 and Vil-Cre:NDI1 n=5. Genes with p-values < 0.05 are shown. (**b**) *NDI1* normalized gene expression in jejunum and liver of the same Vil-Cre:NDI1 mice one hour after oral gavage of vehicle; n=5 (q = 4.46 x 10^-59^, Wald test). All mice were male, 9-12 weeks of age, and fed a standard diet.

**Extended Data Figure 3.**
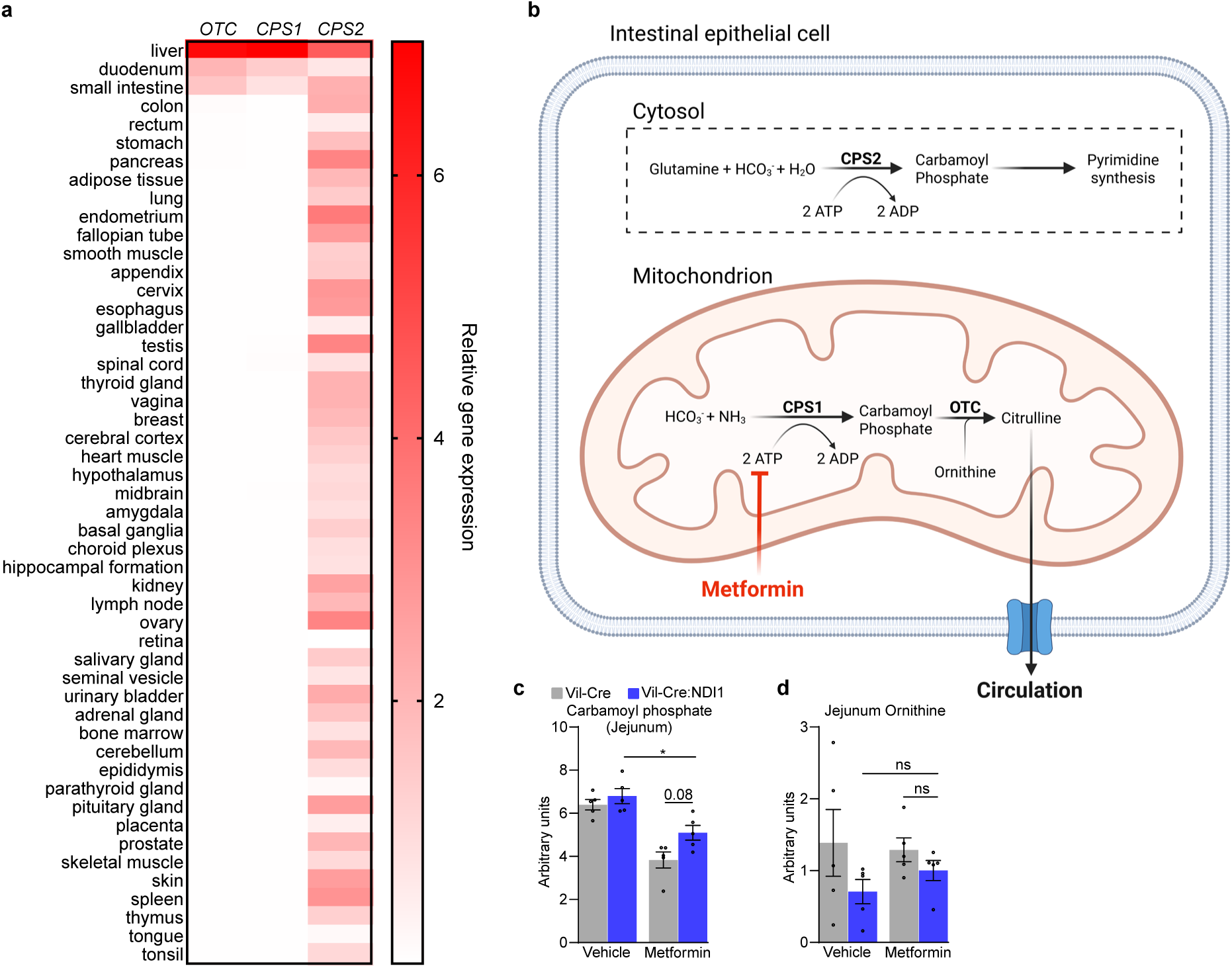
Circulating citrulline is synthesized in the mitochondria of intestinal epithelial cells. (**a**) Relative tissue-specific gene expression profiles for the enzymes that synthesize citrulline (OTC), mitochondrial carbamoyl phosphate (CPS1) and cytosolic carbamoyl phosphate (CPS2/CAD) in humans. Data retrieved 5/5/2025 from consensus RNA expression per gene based on transcriptomic profiles from The Human Protein Atlas and GTEx (https://www.proteinatlas.org/humanproteome/tissue/data#consensus_tissues_rna)^113–115^. (**b**) Schematic illustrating citrulline synthesis in the mitochondria of intestinal epithelial cells. Created with BioRender.com. (**c**) Relative carbamoyl phosphate levels in jejunum one hour after oral administration of vehicle (water) or metformin (200 mg kg^-1^) in overnight-fasted mice; n=5 per condition. (**d**) Relative ornithine levels in jejunum one hour after oral administration of vehicle (water) or metformin (200 mg kg^-1^) in overnight-fasted mice; n=5 per condition. For (c) and (d), data are presented as mean ± SEM. Statistical significance for (c) and (d) was determined by Two-way ANOVA with Bonferroni’s correction for multiple comparisons. *P<0.05.

**Extended Data Figure 4.**
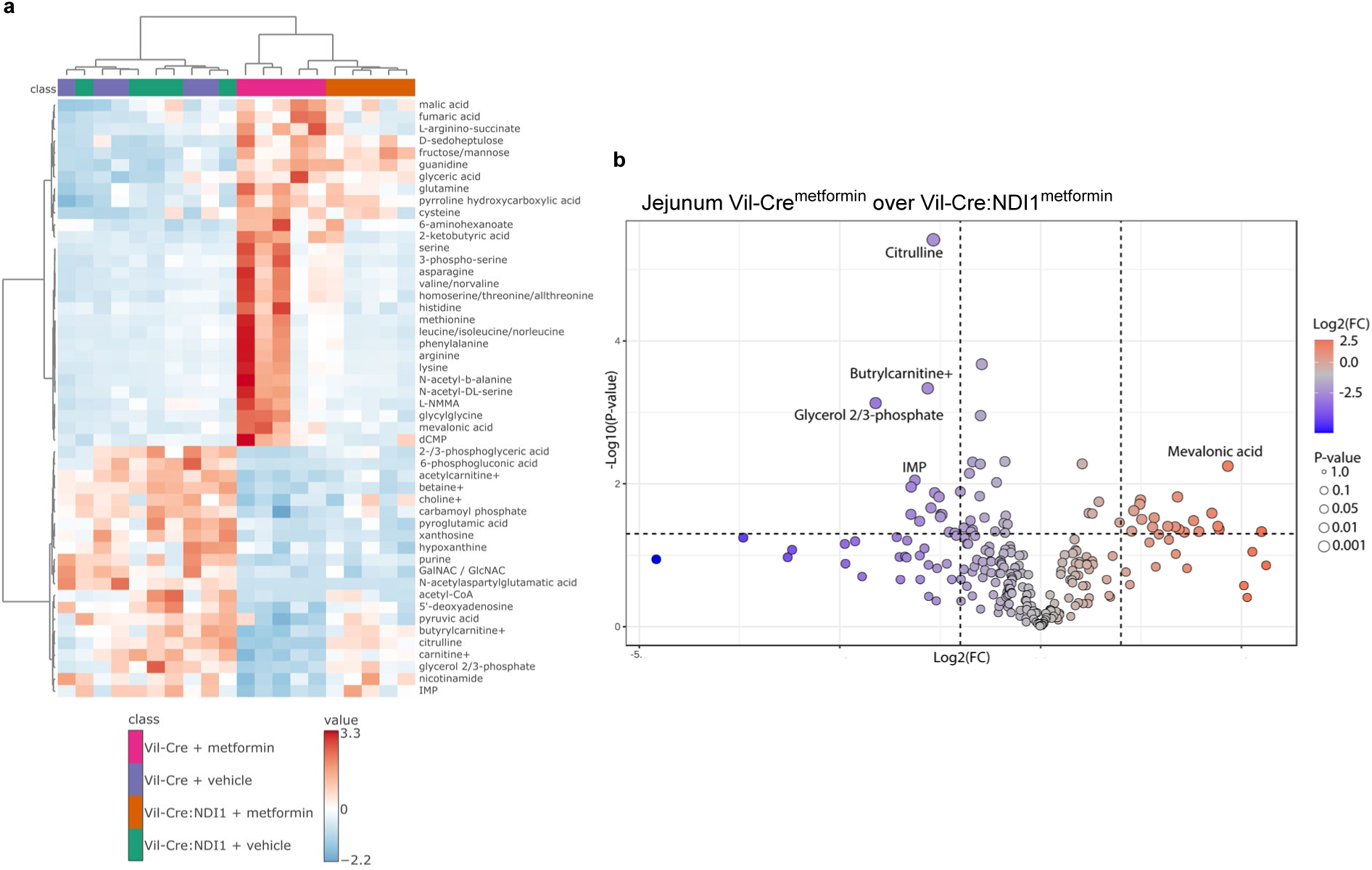
Metformin inhibits mitochondrial complex I to alter the intestinal metabolome. (**a**) Metabolomics heatmap of jejunum one hour after orally administered vehicle (water) or metformin (200 mg kg^-1^) in overnight fasted mice; n=5 per condition. (**b**) Differentially abundant metabolites between Vil-Cre and Vil-Cre:NDI1 mice treated with metformin from (a). The top five metabolites are labeled; n=5 per condition. All mice were male, 9-12 weeks of age, and fed a standard diet.

**Extended Data Figure 5.**
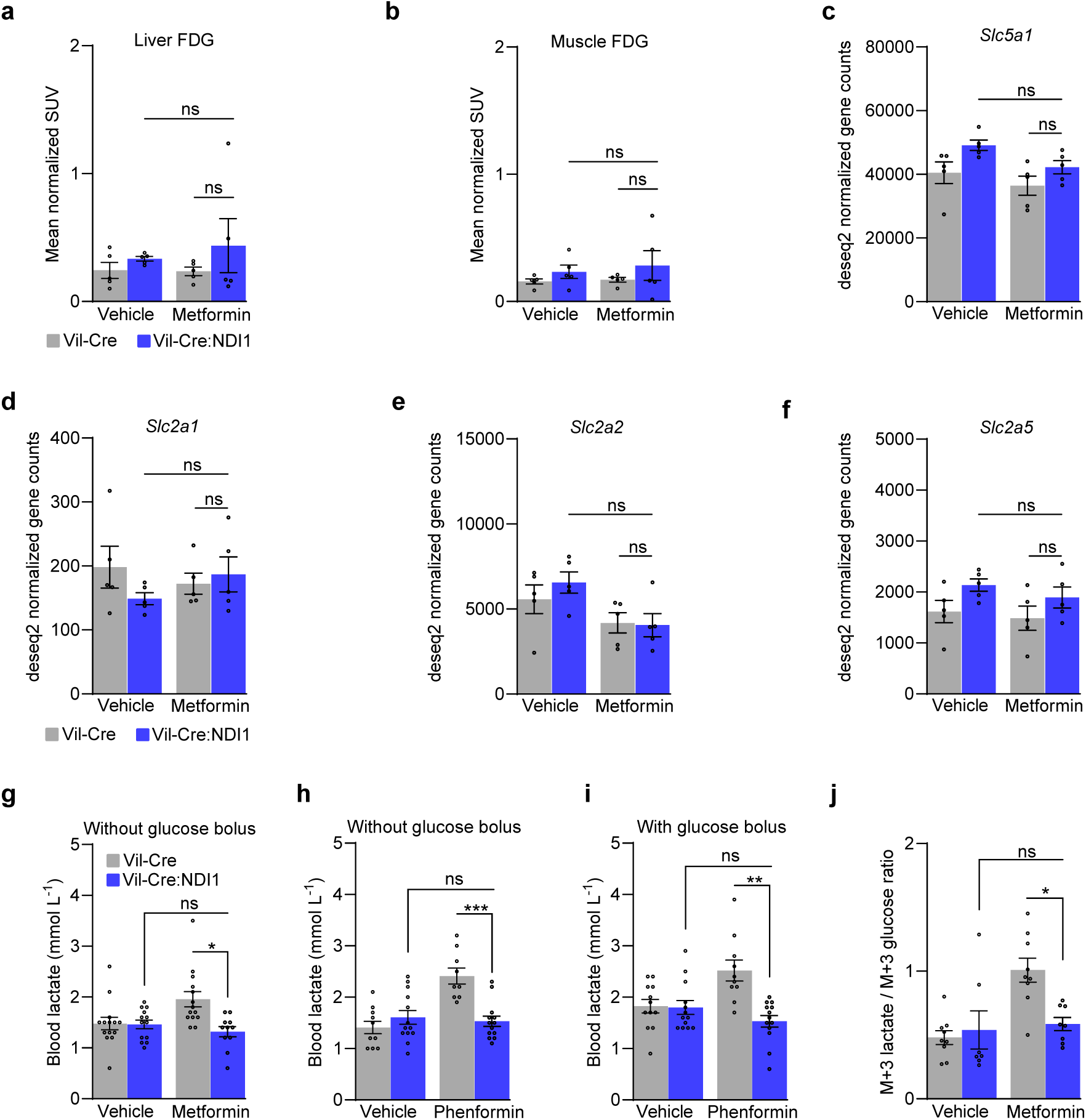
Biguanides inhibit mitochondrial complex I in intestinal epithelium to drive glycolysis. (**a**) Liver FDG accumulation; n=5 per condition. (**b**) Hindlimb muscle FDG accumulation; n=5 per condition. (**c**) Expression of *Slc5a1* (SGLT1) in the jejunum; n=5 per condition. (**d**) Expression of *Slc2a1* (GLUT1) in the jejunum; n=5 per condition. (**e**) Expression of *Slc2a2* (GLUT2) in the jejunum; n=5 per condition. (**f**) Expression of *Slc2a5* (GLUT5) in the jejunum; n=5 per condition. (**g**) Blood lactate one hour after oral administration of vehicle (water) or metformin (200 mg kg^-1^) to overnight-fasted mice; Vil-Cre^vehicle^ n=14, Vil-Cre:NDI1^vehicle^ n=13, Vil-Cre^metformin^ n=14, Vil-Cre:NDI1^metformin^ n=11. (**h**) Blood lactate one hour after oral administration of vehicle (water) or phenformin (100 mg kg^-1^) to overnight-fasted mice; Vil-Cre^vehicle^ n=10, Vil-Cre:NDI1^vehicle^ n=13, Vil-Cre^phenformin^ n=9, Vil-Cre:NDI1^phenformin^ n=13. (**i**) Blood lactate one hour after oral administration of vehicle (water) or phenformin (100 mg kg^-1^) to overnight-fasted mice. An oral bolus of glucose (2 g kg^-1^) was delivered 30 minutes after vehicle/phenformin administration; Vil-Cre^vehicle^ n=12, Vil-Cre:NDI1^vehicle^ n=13, Vil-Cre^phenformin^ n=10, Vil-Cre:NDI1^phenformin^ n=13. (**j**) Plasma m+3 lactate / m+6 glucose ratio of overnight fasted mice one hour after oral administration of vehicle (water) or metformin (200 mg kg^-1^). An oral bolus of U-^13^C_6_-glucose (2 g kg^-1^) was delivered 30 minutes after vehicle/metformin administration; Vil-Cre^vehicle^ n=9, Vil-Cre:NDI1^vehicle^ n=7, Vil-Cre^metformin^ n=9, Vil-Cre:NDI1^metformin^ n=8. All mice were male, 8-12 weeks of age, and fed a standard diet. Data are presented as mean ± SEM. Statistical significance was determined by Two-way ANOVA with Bonferroni’s correction for multiple comparisons. *P<0.05, **P<0.01, ***P<0.001.

**Extended Data Figure 6.**
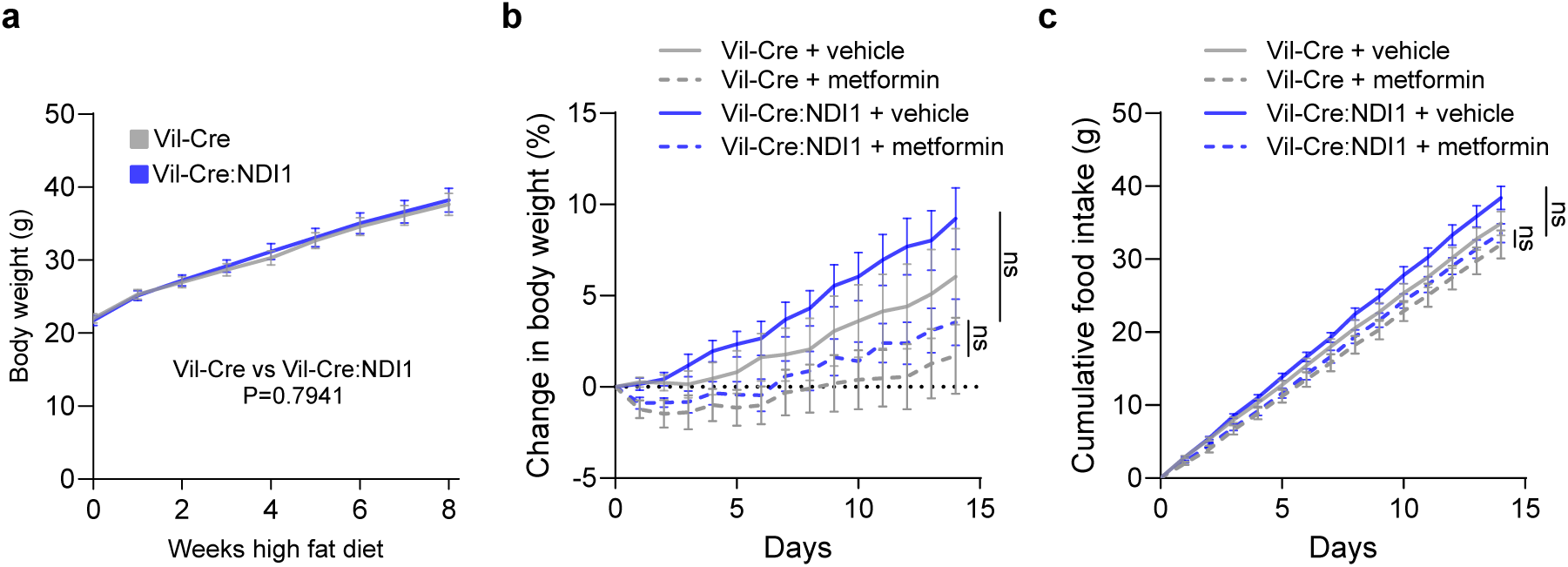
Metformin modestly slows weight gain on high-fat diet. (**a**) Weight gain of Vil-Cre and Vil-Cre: NDI1 mice on a high-fat diet (60% lard) starting at 8 weeks of age; Vil-Cre n=12, Vil-Cre:NDI1 n=17. (**b**) Percent body weight gain of Vil-Cre and Vil-Cre:NDI1 mice on a high-fat diet treated daily with vehicle or metformin (200 mg kg^-1^) by oral gavage; Vil-Cre^vehicle^ n=9, Vil-Cre:NDI1^vehicle^ n=11, Vil-Cre^metformin^ n=9, Vil-Cre:NDI1^metformin^ n=11. (**c**) Cumulative food intake of Vil-Cre and Vil-Cre:NDI1 mice on a high-fat diet (60% lard) treated daily with vehicle or metformin (200 mg kg^-1^) by oral gavage; Vil-Cre^vehicle^ n=7, Vil-Cre:NDI1^vehicle^ n=11, Vil-Cre^metformin^ n=8, Vil-Cre:NDI1^metformin^ n=10. All mice were male and started on high-fat diet at 8 weeks of age. Statistical significance was determined by t-test (a) or Two-way ANOVA with Bonferroni’s correction for multiple comparisons (b and c) at the final time point for each experiment.

**Extended Data Figure 7.**
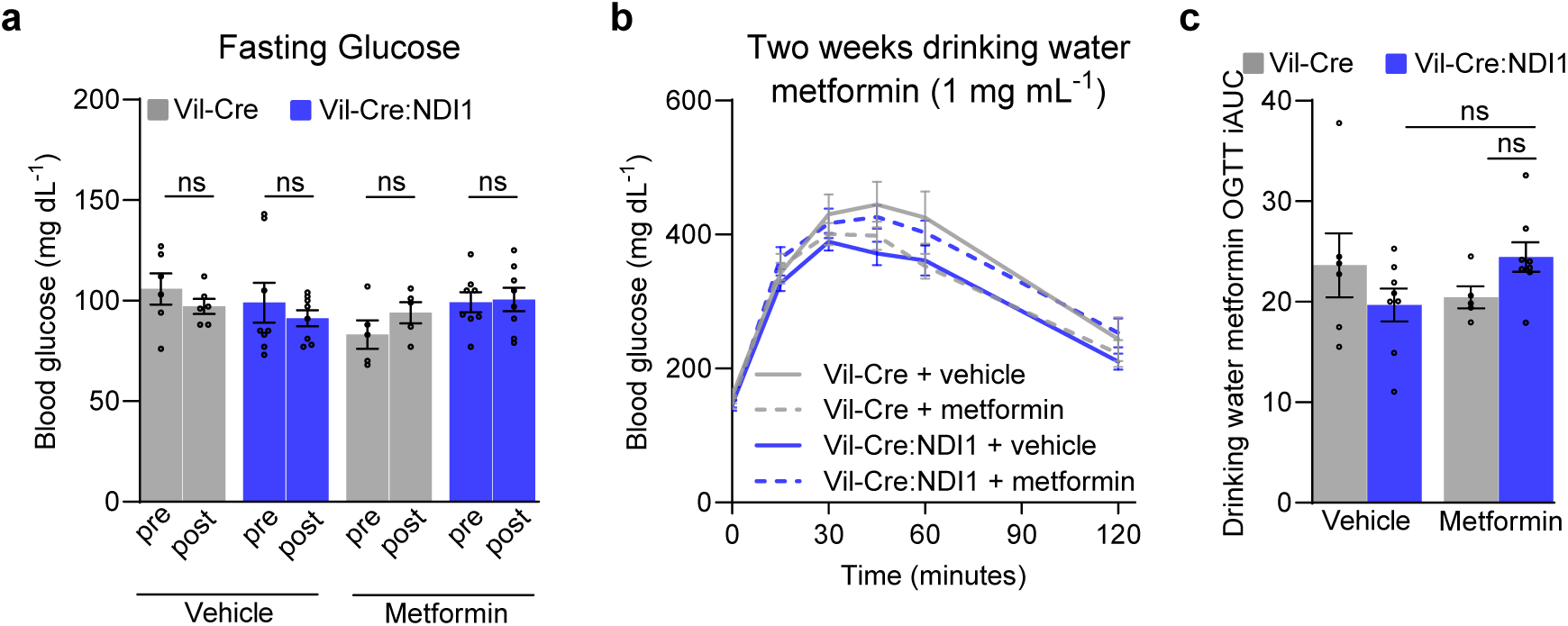
Low-dose metformin administered in drinking water does not lower blood glucose. (**a**) Fasting glucose before and after two weeks of metformin (1 mg mL⁻¹) in drinking water; Vil-Cre^vehicle^ n=6, Vil-Cre:NDI1^vehicle^ n=8, Vil-Cre^metformin^ n=5, Vil-Cre:NDI1^metformin^ n=8. (**b**) Oral glucose tolerance test of diet-induced obese mice treated with metformin (1 mg mL⁻¹) in drinking water for two weeks. Mice were fasted overnight, followed by oral administration of glucose (2 g kg^-1^). (**c**) Incremental area under the curve of (b); Vil-Cre^vehicle^ n=6, Vil-Cre:NDI1^vehicle^ n=8, Vil-Cre^metformin^ n=5, Vil-Cre:NDI1^metformin^ n=8. All mice were fed a HFD for 8 weeks prior to initiation of metformin drinking water. Statistical significance was determined by Two-way ANOVA with Bonferroni’s correction for multiple comparisons (c) or repeated measures Two-way ANOVA with Bonferroni’s correction for multiple comparisons (a).

**Extended Data Figure 8.**
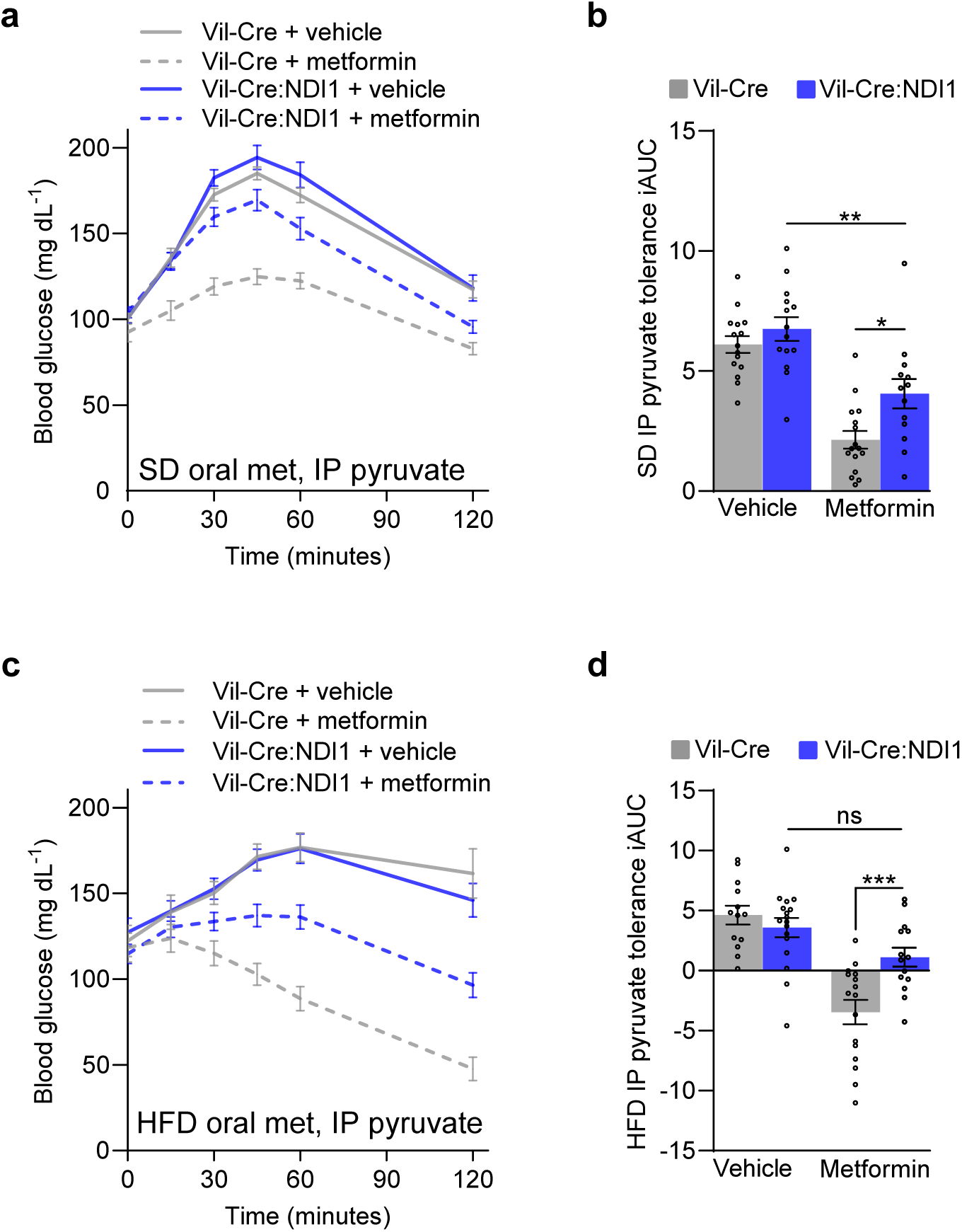
Intestinal mitochondrial complex I inhibition is necessary for metformin to improve intraperitoneal pyruvate tolerance. (**a**) Pyruvate tolerance test in which 200 mg kg^-1^ metformin was delivered by oral gavage 30 minutes before the intraperitoneal injection of pyruvate (2 g kg^-1^) in overnight-fasted mice on standard diet. (**b**) Incremental area under the curve of (a); Vil-Cre^vehicle^ n=15, Vil-Cre:NDI1^vehicle^ n=14, Vil-Cre^metformin^ n=16, Vil-Cre:NDI1^metformin^ n=13. (**c**) Pyruvate tolerance test in which 200 mg kg^-1^ metformin was delivered by oral gavage 30 minutes before the intraperitoneal injection of pyruvate (2 g kg^-1^) in overnight-fasted mice on high-fat diet. (**d**) Incremental area under the curve of (c); Vil-Cre^vehicle^ n=13, Vil-Cre:NDI1^vehicle^ n=17, Vil-Cre^metformin^ n=16, Vil-Cre:NDI1^metformin^ n=14. SD = standard diet; HFD = high-fat diet (60% lard); IP = intraperitoneal; iAUC = incremental area under the curve (arbitrary units). All mice were male. For SD-fed mice, pyruvate tolerance tests were performed on 7–10-week-old animals. For HFD-fed mice, HFD was started at 8 weeks of age and pyruvate tolerance tests were performed after 8-10 weeks of HFD feeding. Data are presented as mean ± SEM. Statistical significance was determined by Two-way ANOVA with Bonferroni’s correction for multiple comparisons. *P<0.05, **P<0.01, ***P<0.001.

**Extended Data Figure 9.**
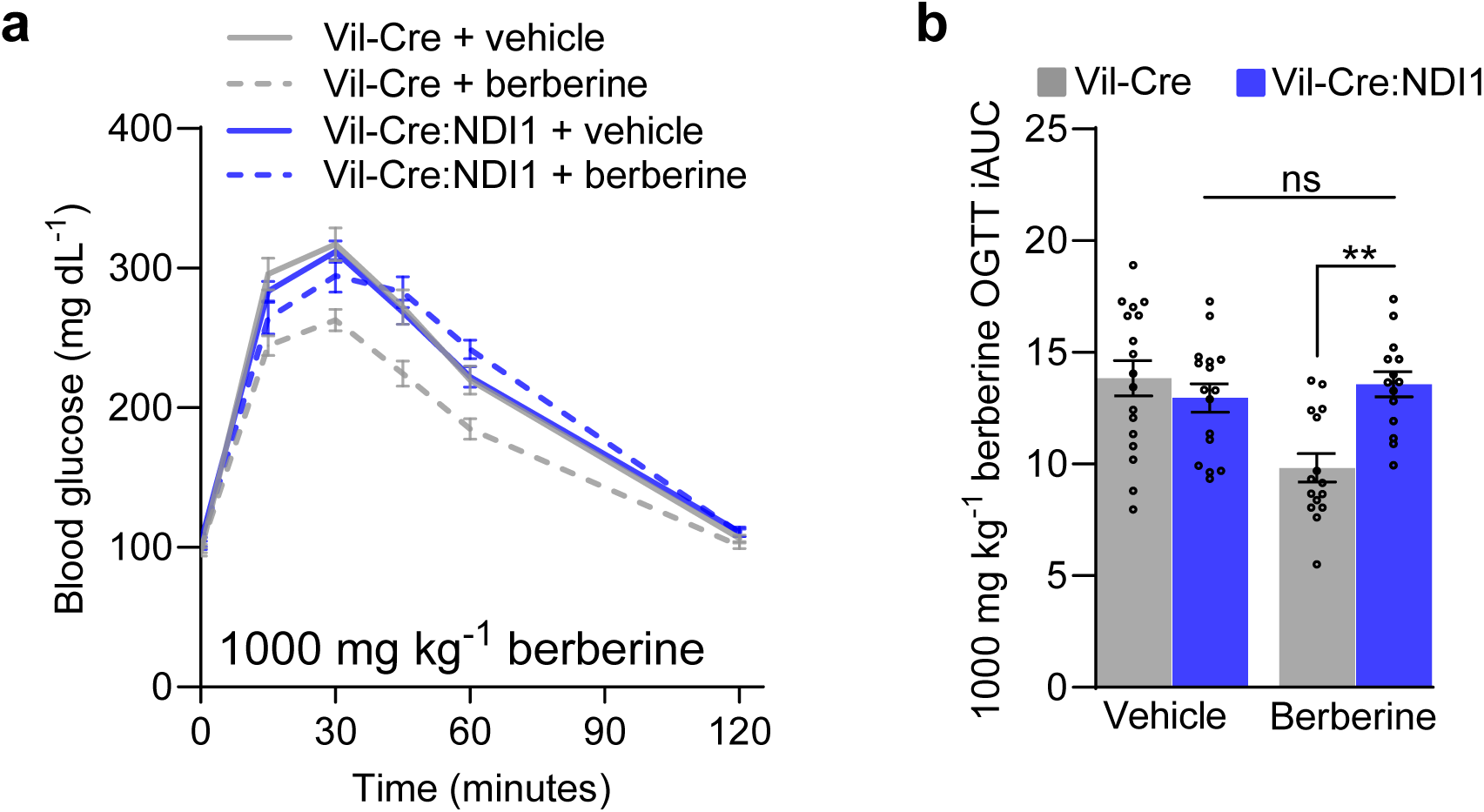
Berberine inhibits mitochondrial complex I in intestinal epithelium to improve glucose tolerance. (**a**) Oral glucose tolerance test with 1000 mg kg^-1^ berberine delivered by oral gavage 30 minutes before the oral gavage of glucose (2 g kg^-1^) in overnight-fasted mice on standard diet. (**b**) Incremental area under the curve of (c); Vil-Cre^vehicle^ n=17, Vil-Cre:NDI1^vehicle^ n=16, Vil-Cre^berberine^ n=15, Vil-Cre:NDI1^berberine^ n=14. Data are presented as mean ± SEM. All mice were male and 7-10 weeks of age. OGTT = oral glucose tolerance test, iAUC = incremental area under the curve (arbitrary units). Statistical significance was determined by Two-way ANOVA with Bonferroni’s correction for multiple comparisons. **P<0.01.

**Extended Data Table 1.**
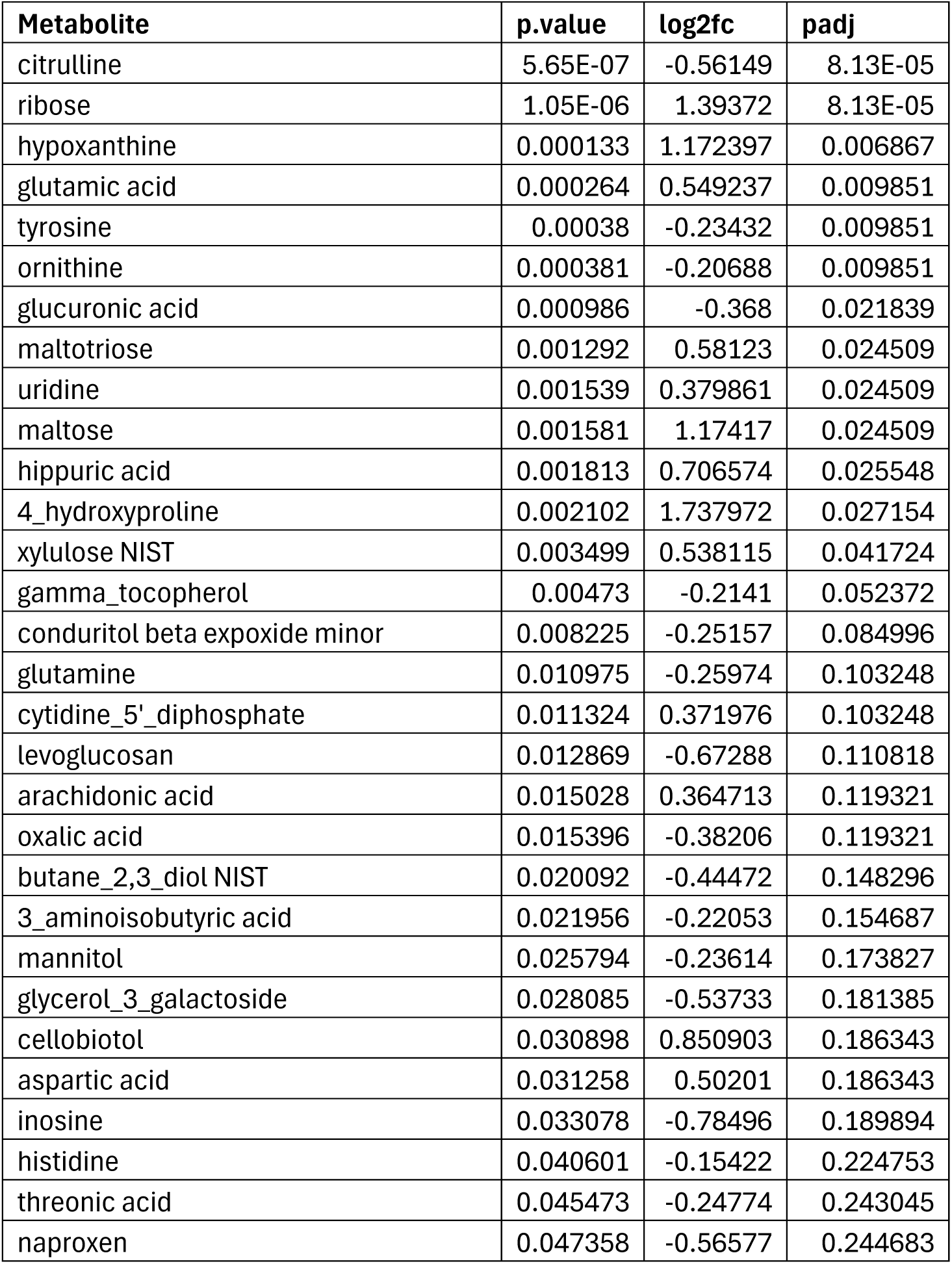
Differentially abundant plasma metabolites before and after metformin treatment in the Rotroff et al. cohort. Metabolites with a raw p-value of less than 0.05 are shown.

| Metabolite | p.value | log2fc | padj |
| --- | --- | --- | --- |
| citrulline | 5.65E-07 | -0.56149 | 8.13E-05 |
| ribose | 1.05E-06 | 1.39372 | 8.13E-05 |
| hypoxanthine | 0.000133 | 1.172397 | 0.006867 |
| glutamic acid | 0.000264 | 0.549237 | 0.009851 |
| tyrosine | 0.00038 | -0.23432 | 0.009851 |
| ornithine | 0.000381 | -0.20688 | 0.009851 |
| glucuronic acid | 0.000986 | -0.368 | 0.021839 |
| maltotriose | 0.001292 | 0.58123 | 0.024509 |
| uridine | 0.001539 | 0.379861 | 0.024509 |
| maltose | 0.001581 | 1.17417 | 0.024509 |
| hippuric acid | 0.001813 | 0.706574 | 0.025548 |
| 4_hydroxyproline | 0.002102 | 1.737972 | 0.027154 |
| xylulose NIST | 0.003499 | 0.538115 | 0.041724 |
| gamma_tocopherol | 0.00473 | -0.2141 | 0.052372 |
| conduritol beta expoxide minor | 0.008225 | -0.25157 | 0.084996 |
| glutamine | 0.010975 | -0.25974 | 0.103248 |
| cytidine_5'_diphosphate | 0.011324 | 0.371976 | 0.103248 |
| levoglucosan | 0.012869 | -0.67288 | 0.110818 |
| arachidonic acid | 0.015028 | 0.364713 | 0.119321 |
| oxalic acid | 0.015396 | -0.38206 | 0.119321 |
| butane_2,3_diol NIST | 0.020092 | -0.44472 | 0.148296 |
| 3_aminoisobutyric acid | 0.021956 | -0.22053 | 0.154687 |
| mannitol | 0.025794 | -0.23614 | 0.173827 |
| glycerol_3_galactoside | 0.028085 | -0.53733 | 0.181385 |
| cellobiotol | 0.030898 | 0.850903 | 0.186343 |
| aspartic acid | 0.031258 | 0.50201 | 0.186343 |
| inosine | 0.033078 | -0.78496 | 0.189894 |
| histidine | 0.040601 | -0.15422 | 0.224753 |
| threonic acid | 0.045473 | -0.24774 | 0.243045 |
| naproxen | 0.047358 | -0.56577 | 0.244683 |

**Extended Data Table 2.**
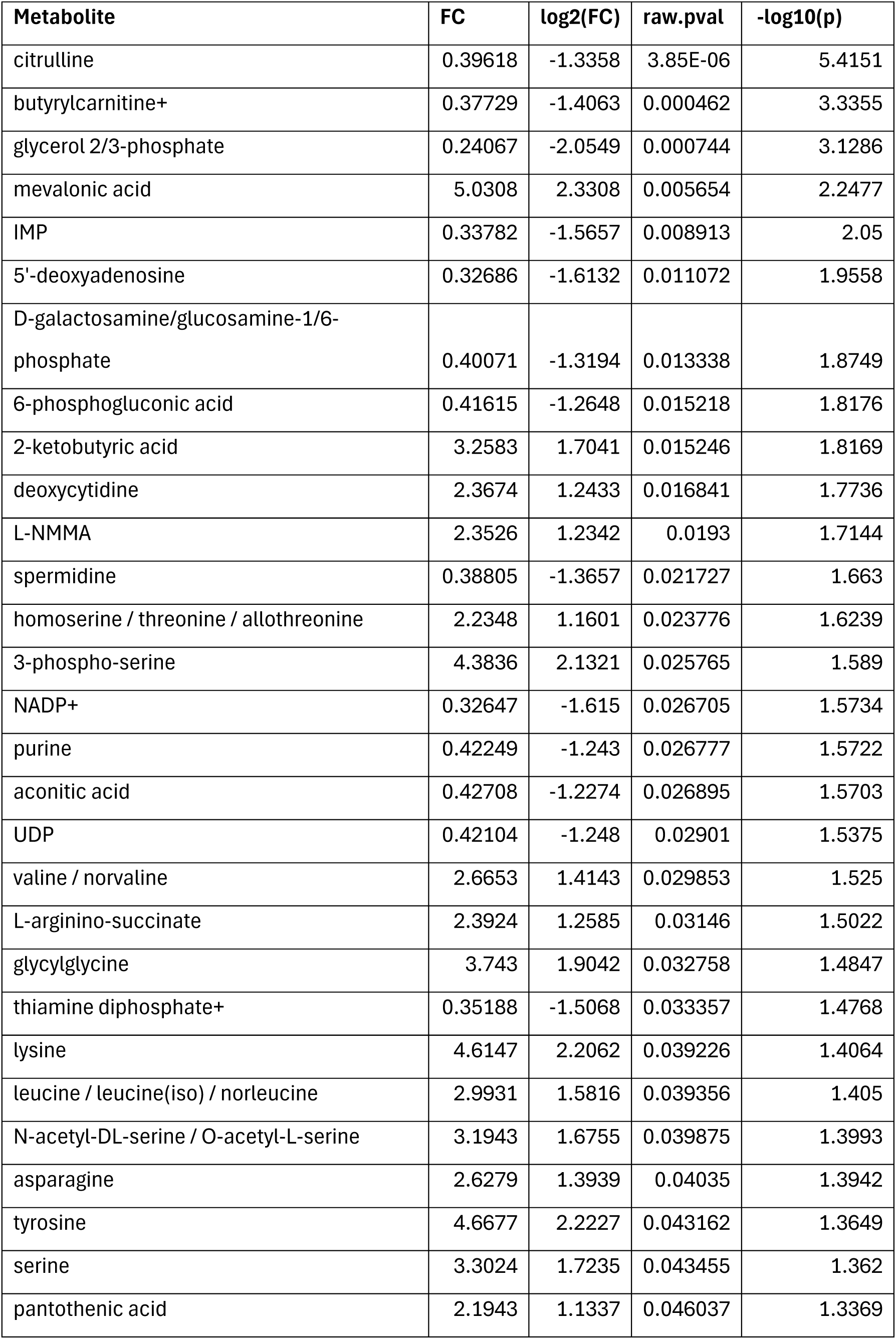
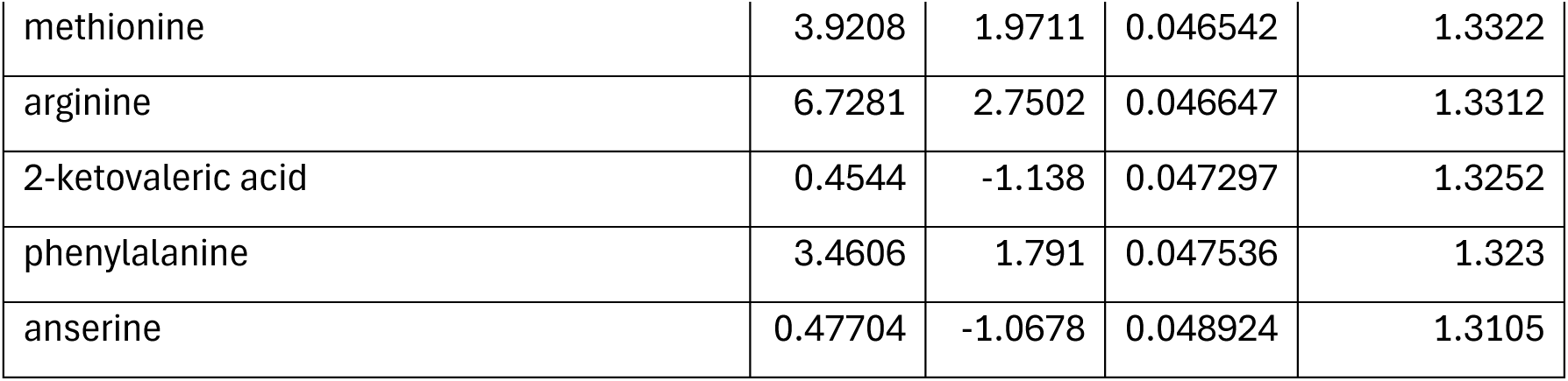
Differentially abundant metabolites between VilCre^metformin^ and VilCre:NDI1^metformin^ jejunum. Overnight-fasted mice were oral gavaged with metformin (200 mg kg^-1^), and jejunum was harvested one hour later. Metabolites with a raw p-value of less than 0.05 are shown.

| Metabolite | FC | log2(FC) | raw.pval | -log10(p) |
| --- | --- | --- | --- | --- |
| citrulline | 0.39618 | -1.3358 | 3.85E-06 | 5.4151 |
| butyrylcarnitine+ | 0.37729 | -1.4063 | 0.000462 | 3.3355 |
| glycerol 2/3-phosphate | 0.24067 | -2.0549 | 0.000744 | 3.1286 |
| mevalonic acid | 5.0308 | 2.3308 | 0.005654 | 2.2477 |
| IMP | 0.33782 | -1.5657 | 0.008913 | 2.05 |
| 5'-deoxyadenosine | 0.32686 | -1.6132 | 0.011072 | 1.9558 |
| D-galactosamine/glucosamine-1/6-phosphate | 0.40071 | -1.3194 | 0.013338 | 1.8749 |
| 6-phosphogluconic acid | 0.41615 | -1.2648 | 0.015218 | 1.8176 |
| 2-ketobutyric acid | 3.2583 | 1.7041 | 0.015246 | 1.8169 |
| deoxycytidine | 2.3674 | 1.2433 | 0.016841 | 1.7736 |
| L-NMMA | 2.3526 | 1.2342 | 0.0193 | 1.7144 |
| spermidine | 0.38805 | -1.3657 | 0.021727 | 1.663 |
| homoserine / threonine / allothreonine | 2.2348 | 1.1601 | 0.023776 | 1.6239 |
| 3-phospho-serine | 4.3836 | 2.1321 | 0.025765 | 1.589 |
| NADP+ | 0.32647 | -1.615 | 0.026705 | 1.5734 |
| purine | 0.42249 | -1.243 | 0.026777 | 1.5722 |
| aconitic acid | 0.42708 | -1.2274 | 0.026895 | 1.5703 |
| UDP | 0.42104 | -1.248 | 0.02901 | 1.5375 |
| valine / norvaline | 2.6653 | 1.4143 | 0.029853 | 1.525 |
| L-arginino-succinate | 2.3924 | 1.2585 | 0.03146 | 1.5022 |
| glycylglycine | 3.743 | 1.9042 | 0.032758 | 1.4847 |
| thiamine diphosphate+ | 0.35188 | -1.5068 | 0.033357 | 1.4768 |
| lysine | 4.6147 | 2.2062 | 0.039226 | 1.4064 |
| leucine / leucine(iso) / norleucine | 2.9931 | 1.5816 | 0.039356 | 1.405 |
| N-acetyl-DL-serine / O-acetyl-L-serine | 3.1943 | 1.6755 | 0.039875 | 1.3993 |
| asparagine | 2.6279 | 1.3939 | 0.04035 | 1.3942 |
| tyrosine | 4.6677 | 2.2227 | 0.043162 | 1.3649 |
| serine | 3.3024 | 1.7235 | 0.043455 | 1.362 |
| pantothenic acid | 2.1943 | 1.1337 | 0.046037 | 1.3369 |
| methionine | 3.9208 | 1.9711 | 0.046542 | 1.3322 |
| arginine | 6.7281 | 2.7502 | 0.046647 | 1.3312 |
| 2-ketovaleric acid | 0.4544 | -1.138 | 0.047297 | 1.3252 |
| phenylalanine | 3.4606 | 1.791 | 0.047536 | 1.323 |
| anserine | 0.47704 | -1.0678 | 0.048924 | 1.3105 |

